# The Western Lake Erie Culture Collection: A promising resource for evaluating the physiological and genetic diversity of *Microcystis* and its associated microbiome

**DOI:** 10.1101/2022.10.21.513177

**Authors:** Colleen E. Yancey, E. Anders Kiledal, Vincent J. Denef, Reagan M. Errera, Jacob T. Evans, Lauren Hart, Dragan Isailovic, William James, Jenan K. Kharbush, Jeffrey A Kimbrel, Wei Li, Xavier Mayali, Helena Nitschky, Catherine Polik, McKenzie A Powers, Sanduni H. Premathilaka, Nicole Rappuhn, Laura A. Reitz, Sara R. Rivera, Claire C. Zwiers, Gregory J. Dick

**Author notes:** Corresponding Author: Dr. Gregory J. Dick, 1100 North University Ave, Rm. 2014, Ann Arbor, MI, USA.

## Abstract

Cyanobacteria harmful algal blooms (cyanoHABs) dominated by *Microcystis* spp. have significant public health and economic implications in freshwater bodies around the world. These blooms are capable of producing a variety of cyanotoxins, including microcystins, that affect fishing and tourism industries, human and environmental health, and access to drinking water. In this study, we isolated and sequenced the genomes of 21 unialgal *Microcystis* cultures collected from western Lake Erie between 2017-2019. While some cultures isolated in different years have a high degree of genetic similarity (Average Nucleotide Identity >99%), genomic data shows that these cultures also represent much of the breadth of known *Microcystis* diversity in natural populations. Only 5 isolates contained all the genes required for microcystin synthesis while 2 isolates contained a previously described partial *mcy* operon. Microcystin production within cultures was also assessed using Enzyme-Linked Immunosorbent Assay (ELISA) and supported genomic results with high concentrations (up to 900 μg L-1) in cultures with complete *mcy* operons and no or low toxin detected otherwise. These xenic cultures also contained a substantial diversity of bacteria associated with *Microcystis*, which has become increasingly recognized as an essential component of cyanoHAB community dynamics. These results highlight the genomic diversity among *Microcystis* strains and associated bacteria in Lake Erie, and their potential impacts on bloom development, toxin production, and toxin degradation. This collection significantly increases the availability of environmentally relevant *Microcystis* strains from temperate North America, which is changing rapidly due to climate change.

**Highlights:**

- Twenty one xenic *Microcystis* cultures were isolated from western Lake Erie and capture the diversity of *Microcystis* strains observed in natural populations as well as their associated bacteria
- *Microcystis* strains show variability in core and accessory gene content, and genetically similar strains produce varying concentrations and congeners of microcystins
- This collection is a valuable resource for studying strain diversity and interactions between *Microcystis* and associated bacteria
- Our collection increases the availability of environmentally relevant strains from temperate North America, which is historically underrepresented in culture collections.

## Introduction

The Laurentian Great Lakes are increasingly threatened by cyanobacteria harmful algal blooms (cyanoHABs) capable of producing cyanotoxins that are harmful to human and environmental health (Binding et al., 2020; McKindles et al., 2020; Miller et al., 2017). Of particular concern is western Lake Erie (WLE), which experiences intense cyanoHABs dominated by the microcystin (MC) producing cyanobacterium *Microcystis* spp. (Berry et al., 2017; Bridgeman et al., 2013; Rinta-Kanto et al., 2005; Steffen et al., 2017; Watson et al., 2016). These annual blooms begin midsummer and can persist through late fall, threatening drinking water supplies to both Canada and the United States. Blooms can also cause hypoxia, fish kills, and other ecosystem disruptions (Berry et al., 2017; Harke et al., 2016; Huisman et al., 2018; Steffen et al., 2017; Watson et al., 2016). CyanoHABs have substantial economic ramifications as well: a single bloom in WLE during August 2014 led to a $65 million loss in the United States (Bingham et al., 2015), and blooms in WLE will cost Canada an estimated $5.3 billion over the next 30 years if left unmanaged (Smith et al., 2019). The threat to potable water access is of greatest concern and became a reality during the 2014 cyanoHAB, resulting in a “do not drink” advisory in the city of Toledo, Ohio, USA due to high levels of MC in treated drinking water supplies (Steffen et al., 2017).

*Microcystis*-dominated cyanoHABs are expected to increase in intensity and frequency both globally and within WLE as a result of intensifying climate change (Griffith and Gobler, 2020; Ho et al., 2019; Huisman et al., 2018; Watson et al., 2016). Toxin content per cell within blooms is also expected to increase due to climate change associated increases in atmospheric *pCO2* (Sandrini et al., 2015) and the continuation of unchecked nitrogen (N) loading (Hellweger et al., 2022; Wagner et al., 2019). As climate change continues to alter freshwater systems, it is critical to understand the effects of this change to better inform future management strategies.

The diversity of *Microcystis* strains and their interactions with associated bacteria in the phycosphere are potentially important controls on toxin content and concentration (Dick et al., 2021). Natural phytoplankton assemblages are known to contain multiple strains of *Microcystis*, some capable of producing MC while others are not (Rinta-Kanto et al., 2005, 2009; Yancey et al., 2022a). Further, the *mcy* operon, which encodes the synthesis of MCs (Tillett et al., 2000), exhibits large sequence diversity between strains which may contribute to differential congener production (Mikalsen et al., 2003; Tooming-Klunderud et al., 2008; Yancey et al., 2022a). For example, the drivers of the emergence of MC-LA in Lake Erie (Birbeck et al., 2019), a congener typically seen on the West Coast of the United States, remains elusive and may be the result of changing strain composition and/or environmental gradients. Another dimension of *Microcystis* diversity is the microbiome of bacteria that are physically associated with *Microcystis* colonies. These microbiomes are *Microcystis* strain-specific (Smith et al., 2021) and likely engage in mutualistic interactions with *Microcystis* (Smith et al., 2022a, 2022b). While such interactions influence toxin production by other algae (Bates et al., 1995), the effects on toxin production by *Microcystis* are largely unknown. Thus, it is imperative that we understand the intricate dynamics of cyanoHABs including *Microcystis* strain diversity and interactions within the cyanoHAB phycosphere in order to understand how toxin production and bloom progression may be impacted by anthropogenic forcings.

The genus *Microcystis* is well known for its complex genomes, rich with mobile elements (Harke et al., 2016), low synteny among strains (Humbert et al., 2013), and a large pangenome (Cao et al., 2022; Meyer et al., 2017). *Microcystis* shows substantial strain-level diversity (Harke et al., 2016; Humbert et al., 2013), with extensive heterogeneity of genetic and phenotypic characteristics (Dick et al., 2021). High rates of horizontal gene transfer among *Microcystis* strains (Cao et al., 2022; Pérez-Carrascal et al., 2019) complicate subspecies delineation and may contribute to their environmental success and adaptation to variable environments (Cao et al., 2022; Dick et al., 2021). Their large accessory genomes and potentially global pangenome (Meyer et al., 2017) harbor numerous genes of known and unknown function, many of which may encode toxins (Pérez-Carrascal et al., 2019) or other traits that provide *Microcystis* a competitive advantage in natural communities. While functional groups and key traits within this genus have yet to be fully defined, it is critical to understand how intraspecies *Microcystis* diversity affects the toxicity, persistence, and dominance of *Microcystis* in blooms around the world (Dick et al., 2021).

Several methods are commonly used to investigate cyanoHAB dynamics including long term field monitoring and sampling, and controlled culture experiments in mesocosm and laboratory settings. Field samples provide direct insight into entire cyanoHAB communities, but the complexity of whole communities can complicate the assignment of particular functions to specific taxa. Axenic cultures enable insights into the genetic and physiological traits of specific *Microcystis* strains (Dick et al., 2021; Harke et al., 2016; Humbert et al., 2013; Pérez-Carrascal et al., 2019), but may misrepresent how *Microcystis* functions within a natural community, interacts with its environment, and responds to environmental change, as the associated bacteria are increasingly recognized to be essential for cyanoHAB dynamics (Hoke et al., 2021; Seymour et al., 2017; Smith et al., 2021). Xenic *Microcystis* cultures are valuable as an intermediary between the full complexity of whole communities and the unrealistic simplicity of axenic cultures and can be valuable for studying interactions between *Microcystis* and its associated bacteria. However, there is currently only one publicly available strain of *Microcystis* from WLE: strain LE-3. This strain was isolated over 20 years ago, produces MCs, and is maintained in culture with associated bacteria, (Brittain et al., 2000), but it has lost its ability to form colonies, which are the dominant phenotype in the field, and define the physicochemical habitat of the phycosphere that is packed with heterotrophic bacteria (Smith et al., 2021). Further, LE-3 represents just a single strain, whereas WLE blooms contain multiple diverse strains that play key roles in bloom dynamics (Rinta-Kanto et al., 2005; Yancey et al., 2022a, 2022b).

To address the lack of WLE *Microcystis* isolates in culture collections, we isolated 21 new strains of *Microcystis* from different years, locations, and seasons and conducted initial characterization and genome sequencing. We find that these xenic cultures represent much of the known genetic diversity of *Microcystis* and contain several bacteria known to associate with *Microcystis* in nature. These cultures will serve as a useful resource to better understand the genetic and phenotypic diversity of *Microcystis* and their interactions with associated bacteria.

## 2. Materials and Methods

### 2.1 Collection and Isolation

Xenic, unialgal cultures that contained *Microcystis* were collected for isolation and cultivation during weekly cyanoHAB cruises from various NOAA Great Lakes Environmental Research Laboratory (GLERL) sampling stations within western Lake Erie (Cooperative Institute for Great Lakes Research, n.d.) during September 2017, June and August of 2018, and July and August of 2019 at stations WE2, WE4, WE6, WE8, and WE12 (Cooperative Institute for Great Lakes Research, n.d.; Den Uyl et al., 2022). Further information regarding collection date, location, and isolation technique is outlined in Table 1. Isolation work was performed on either whole water samples or a >100 μm fraction collected using a net tow. Serial dilutions (1x, 10x, 100x, and 1000x) of water samples were plated (see below for media recipes) and incubated at 23°C (room temperature) and 50 μmol photons meter^-2^ second^-1^ on a 12:12 hour light:dark cycle. Colonies identified as *Microcystis* by microscopy were then serially streaked onto agarose from the initial lawn. Once unialgal, cultures continued to be transferred on solid plates and were used to inoculate 0.5 mL liquid cultures. All cultures were cryopreserved in a 5% DMSO solution and stored at −150°C in liquid nitrogen.

**Table 1:** Summary of Culture Isolation and Maintenance.

| Strain group | Culture | Isolation Method | Isolation Media | Collection Date | Isolation Location | *mcy* genotype | Associated bacteria catalase |
| --- | --- | --- | --- | --- | --- | --- | --- |
| 1 | LE17-10 | Solid Dilution | BG11 2N | 9/18/2017 | WE8-1 | Absent | Yes |
| 1 | LE19-196.1 | Solid 1000X Dilution | LE BG11 2N | 8/5/2019 | WE6 | Absent | Yes |
| 1 | LE19-4.1 | Solid 100X Dilution | LE BG11 2N | 7/8/2019 | WE8 | Absent | Yes |
| 2 | LE19-114.1 | Solid 100X Dilution | LE BG11 2N | 7/22/2019 | WE8 | Absent | Yes |
| 3 | LE19-131.1 | Solid 1000X Dilution | BG11 2N | 7/29/2019 | WE8 | Absent | Yes |
| 4 | LE19-55.1 | Solid 100X Dilution | LE WC (1/2X) | 7/15/2019 | WE2 | Absent | Yes |
| 4 | LE19-98.1 | Solid 10X Dilution | BG11 2N | 7/29/2019 | WE4 | Absent | Yes |
| 5 | LE18-13.4 | Solid 10X Dilution | WC (1/2X) | 6/12/2018 | WE2 | Absent | No |
| 6 | LE19-12.2 | Solid 1000X Dilution | LE WC (1/2X) | 7/8/2019 | WE8 | Absent | Yes |
| 6 | LE19-197.1 | Solid 1000X Dilution | BG11 2N | 8/5/2019 | WE8 | Absent | No |
| 7 | LE17-20 | Solid Dilution | BG11 2N | 9/18/2017 | WE8-2 | Absent | Yes |
| 7 | LE19-8.1 | Solid 1000X Dilution | WC (1/2X) | 7/8/2019 | WE8 | Absent | Yes |
| 8 | LE19-84.1 | Solid 100X Dilution | LE WC (1/2X) | 7/29/2019 | WE4 | Complete (B1) | No |
| 9 | LE19-338.1 | Solid 100X Dilution | WC (1/2X) | 8/20/2019 | WE6 | Absent | Yes |
| 9 | LE19-388.1 | Solid 100X Dilution | BG11 (no nitrate) | 7/29/2019 | WE6 | Absent | Yes |
| 10 | LE19-41.2 | Solid 1000X Dilution | LE BG11 2N | 7/15/2019 | WE2 | Complete (C1) | Yes |
| 10 | LE19-59.1 | Solid 1000X Dilution | WC (1/2X) | 7/15/2019 | WE12 | Complete (C1) | Yes |
| 11 | LE19-10.1 | Solid 100X Dilution | BG11 2N | 7/8/2019 | WE8 | Partial (C1) | Yes |
| 11 | LE19-251.1 | Solid 10X Dilution | LE BG11 2N | 8/12/2019 | WE8 | Partial (C1) | Yes |
| 12 | LE18-22.4 | Solid 10X Dilution | WC (1/2X) | 6/25/2018 | WE12 | Complete (C1) | Yes |
| 12 | LE19-195.1 | Solid 100X Dilution | LE BG11 2N | 8/5/2019 | WE6 | Complete (B1) | Yes |

Cultures are maintained on solid plates and in 10 mL liquid media at room temperature and 40 μmol photons meter^-2^ second^-1^ on a 12:12 hour light:dark cycle. Cultures have successfully been maintained at temperatures from 20 to 26°C and light ranges 40 to 600 μmol photons meter^-2^ second^-1^. Acclimation changes of no more than 1°C or 20% μmol photons meter^-2^ second^-1^ each day with weekly transfers into fresh media were required to ensure growth and healthy cultures. Various culture media were used during isolation and maintenance of the culture collection including BG-11 2N (Humbert et al., 2013), which is BG-11 (Allen, 1968) modified to contain 2 mM NaNO_3_ and 10 mM NaHCO_3_, BG11-2N with 0.22 μm filtered Lake Erie water instead of MilliQ water (LE BG11-2N), unmodified BG-11, and half strength Wright’s cryptophyte (WC) medium (Guillard and Lorenzen, 1972) modified according to (Stemberger, 1981) and supplemented with 0.6 μM H_2_O_3_Se. The media in which each culture was isolated is described in Table 1.

### 2.2 DNA Extraction and Sequencing

Cultures grown in BG11-2N with a 12:12 hour diurnal cycle were collected around the 7th day after passage, homogenized, and centrifuged for 5 minutes at 15,000 G twice to collect biomass from 4 mL of dense culture. The DNeasy Blood and Tissue Kit with the QIAshredder adapter (Qiagen, Hilden, Germany) was used to extract DNA according to the protocol found at https://www.protocols.io/view/dna-extraction-from-filters-using-qiagen-dneasy-an-j8nlk61wv5r7/v1. Quan-iT™PicoGreen™ DNA Assay Kit (Eugene, OR, USA) was used to quantify DNA concentrations. Sequencing was completed at the Advanced Genomic Core at the University of Michigan on the NovaSeq (S4) for 300 cycles. The sequencing run was completed with maximum insert possible without compromising read quality.

### 2.3 Metagenomic Analysis

Raw reads were processed with bbduk from JGI supported software package bbtools v.38.84 (https://sourceforge.net/projects/bbmap/, (Bushnell, n.d.)) to remove adapters, quality trim reads, and remove contamination using the Univec reference collection (https://www.ncbi.nlm.nih.gov/tools/vecscreen/univec/). Duplicate reads were removed using clumpify and dedupe (bbtools; see above). Each sample was assembled separately using Megahit v1.2.9 (Li et al., 2015) with the meta-sensitive parameter. Contiguous sequences, or contigs, greater than 1 kb in length, were used to create an Anvi’o database for each sample. *Microcystis* bins and bins of the associated bacteria were generated with Concoct v.1.1.0 (Alneberg et al., 2013) then manually refined with Anvi’o v.7 (Eren et al., 2021, 2015). Bin taxonomy was determined using single copy genes in Anvi’o and additionally evaluated with GTDBtk and the GTDB release 202 database (Chaumeil et al., 2019; Parks et al., 2018). A pangenome analysis of 159 *Microcystis* reference genomes and the 21 genomes obtained here determined 26 single copy genes that were present in each of the 180 genomes. These single copy genes were concatenated and aligned with MUSCLE (Edgar, 2004), and used to produce a tree with fasttree (Price et al., 2009) using the gamma parameter.

For each sample, the relative abundance of organisms was determined by summarizing reads mapped to bins using CoverM v.0.6.1 and the Minimap2 aligner v.2.24 (Li, 2018, p. 2; Woodcroft, n.d.). To screen for un-binned organisms, taxonomic profiles were also produced for each sample using Kraken 2 version 2.1.2 and Bracken version 2.6.1 with a GTDB release 202 database produced with Struo2 (Lu et al., 2017; Wood et al., 2019; Wood and Salzberg, 2014; Youngblut and Ley, 2021). To reduce spurious hits, only taxa with ≥3 minimum-hit-groups and ≥ 150 distinct minimizers were retained; the --report-minimizer-data option was used to obtain these values similar to the unique k-mer counts produced by KrakenUniq (Breitwieser et al., 2018).

Assembled *Microcystis* genomes were screened for various genes including those involved in biosynthesis of secondary metabolites, nitrogen and phosphorus metabolism, carbon concentrating mechanisms (CCM), vitamin synthesis and requirement, and hydrogen peroxide decomposition by catalase. Assembled genomes of associated bacteria were additionally searched for microcystin degrading genes, vitamin B1 and methionine synthesis, and catalase genes. The list of genes queried with NCBI accession numbers can be found in Table S1. BLAST v.2.9.0 (Madden, 2013) was used to align the gene database to the *Microcystis* and associated bacteria isolates. For each genome, a presence/absence matrix was generated and used for further analysis and visualization; genes with >95% identity and >50% alignment length were considered present. Since biosynthetic genes can be short (<1 kb), have high sequence variance, and do not always assemble or bin well (Dick et al., 2021; Humbert et al., 2013; Yancey et al., 2022a), their presence or absence was confirmed by aligning QC’ed reads to the BLAST gene database with the same criteria. CCM genes were analyzed at the read level as well to determine presence/absence. Proteins in the MC degradation pathway were identified using the hmmsearch function in HMMER software package (Version 3.3.2) based on representative custom Hidden Markov model (HMM) profiles including *mlrA-D* genes (Eddy, 2011). A threshold of E-value < 0.001 was used to identify positive hits. To generate the custom profiles, MlrA-D protein sequences available in NCBI were aligned using Clustal Omega (www.ebi.ac.uk/Tools/msa/clustalo/) with default parameters to create seed proteins. HMM profiles were then generated from the alignments using the hmmbuild function in HMMER.

Read mapping was used to deeply evaluate *mcy* operon genotypes present within samples. This was visualized via read mapping plots generated by the plot-blast-frag-cov script found at https://github.com/Geo-omics/scripts. Based on findings previously described (Yancey et al., 2022a), coverage patterns were used to determine presence of the complete (all *mcy* genes present), absent (no *mcy* genes present), or partial (truncated *mcyA*, complete *mcyB* and *mcyC) mcy* genotypes for the *Microcystis* isolates. Variants in the *mcyB* gene, described in a previous study (Mikalsen et al., 2003), were assessed as well.

Additional metabolic functional annotation was done to assess the completeness of pathways using a collection of KEGG orthology terms (searched with kofamscan v1.3.0) or custom HMMs (searched with HMMer v3.3.2). We focused on metabolism for the cycling of substrates, growth hormones and vitamins between *Microcystis* and associated bacteria.

### 2.4 Microcystin Analysis

To determine the overall concentration of MC in cultures after 14 days of growth, Enzyme-Linked Immunosorbent Assays (ELISA) were used. 3-10 mL of liquid culture was filtered through 0.2 um Polycarbonate (PC) membrane filters (Merck Millipore Ltd, Tullgreen, Carrigtwohill, Co Cork, Ireland), until clogged. Samples were stored at −80°C prior to extraction following the EPA Method 546 for detection of total microcystins Method 546 (US EPA, 2019) for particulate microcystin. For comparison, two independent measurements were completed with independently grown cultures. In addition to ELISA assays, MC congener composition was determined using liquid chromatography-mass spectrometry (LC-MS) via methods previously described (Baliu-Rodriguez et al., 2022; Palagama et al., 2020). Congener identification was confirmed via standards for isoforms MC-LR, RR, YR, LA, LW, LF, HilR, LY and D-Asp MC-LR.

## 3. Results

### 3.1 Culture Collection Overview

Isolation of *Microcystis* strains from four locations (Fig. S1) in WLE during 2017-2019 yielded 21 unique, xenic, and *Microcystis* cultures (Table 1) from samples that spanned a variety of environmental conditions (Table S2). The cultures also contain other bacteria, many of which are physically attached to *Microcystis* colonies in the field (Smith et al., 2021). The majority of cultures were isolated from nearshore stations, where cyanobacterial biomass tends to peak in abundance during WLE cyanoHABs (Berry et al., 2017; Bridgeman et al., 2013; Steffen et al., 2017; Watson et al., 2016). Most isolates were collected during July, although some were collected during August, often the time of peak cyanobacterial abundance, and during June (prebloom) and September (post peak) (Table 1).

Currently, all 21 cultures are being maintained at the University of Michigan. These cultures have maintained their ability to form colonies, in contrast to LE-3, which grows unicellularly or in duplets in culture. Variations of the BG11-2N media were the most successful for isolation and acclimation to lab conditions (13/21 cultures) (Table 1). We found that BG-11 2N prepared with milliQ water was sufficient for maintenance due to limitation of collection and storage of Lake Erie water. Unmodified BG-11 and half strength WC media yielded variable success in isolation and maintenance of the WLE culture collection (data not shown, Table 1).

### 3.2 Genome Content and Diversity of WLE Microcystis Isolates

Shotgun metagenome sequencing produced near-complete genomes of *Microcystis* and associated bacteria in the cultures (Table S3). Because other bacteria were present, the shotgun sequence datasets were analyzed as metagenomes. In several cases, the coverage of *Microcystis* genomes was low, around (6-8x), which is below the minimum coverage required for high-quality genome assembly (Dick, 2018), resulting in fragmented assemblies with low N50 (the length of the shortest contig where at least 50% of the assembly is contained in equal or greater length contigs). For example, the *Microcystis* MAG in LE19-196.1 had a reported N50 of 4062 and 1365 contiguous sequences (Table S3). Several of the cultures displayed substantial strain heterogeneity, particularly LE19-12.2, indicating the likely presence of multiple *Microcystis* strains (Table S3). The fraction of sequences mapping to *non-Microcystis* MAGs varied from 0.5% to 86.9% and the number of MAGs retrieved from *non-Microcystis* bacteria varied from one in LE19-84.1 (*Novosphingobium* sp.) to 15 in LE19-196.1.

WLE *Microcystis* isolates are scattered across the phylogenetic tree of all sequenced *Microcystis* genomes (Fig. 1). In some cases, WLE strains cluster tightly with *Microcystis* strains isolated from other locations around the world. For example, LE19-196.1, LE17-10, LE19-114.1 form a tight clade with strains NIES 87, L211 07 and Sj., which were isolated from Japan, the United States, and Japan, respectively. WLE *Microcystis* strains are largely found within groups Mae1, Mae_5, Mfl_ae1, Mflae_ae4, Mae4, and Mae3 (Fig 1). Groups within this tree were previously described (Dick et al., 2021) and are largely congruent with previously described groups (Pérez-Carrascal et al., 2019).

**Figure 1:**
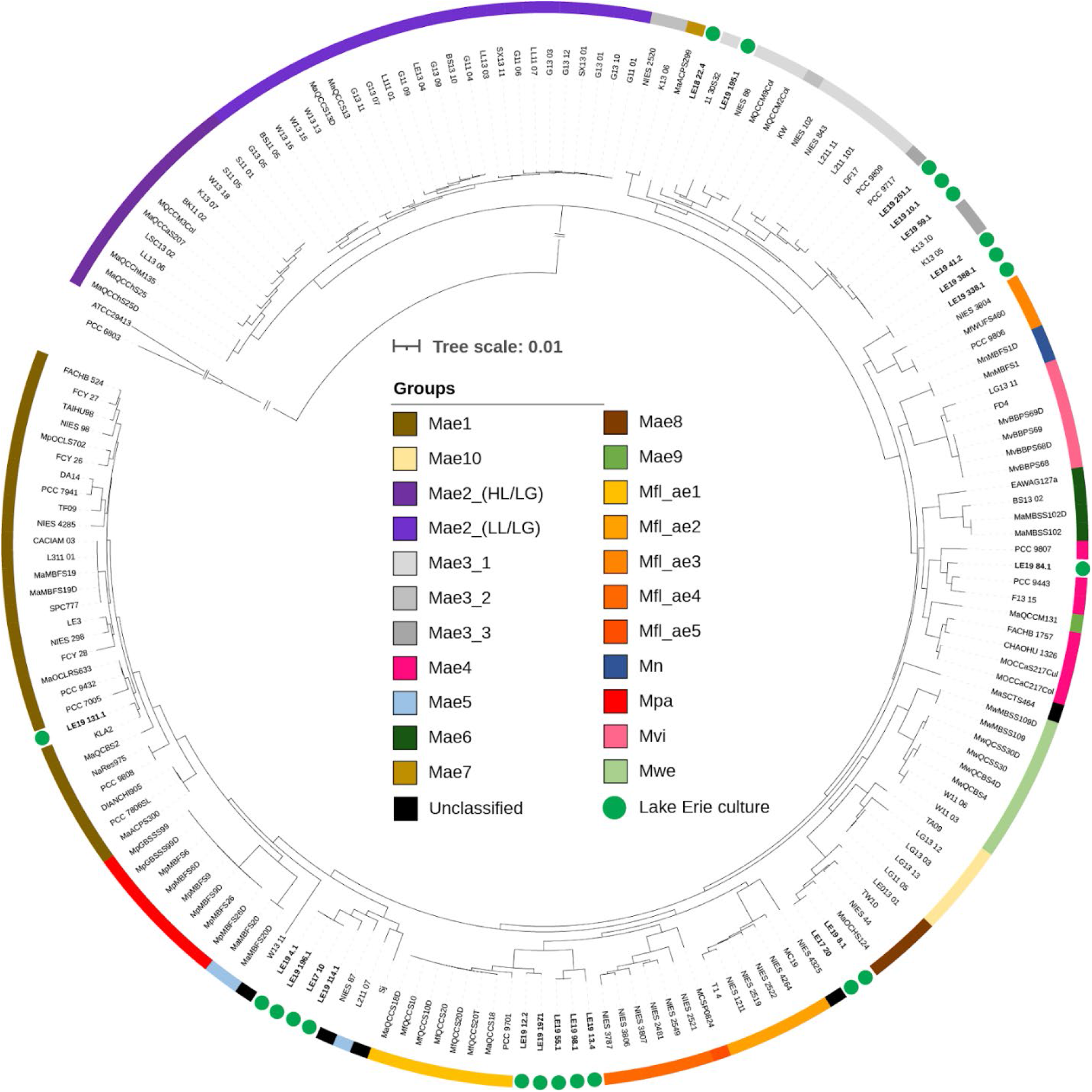
Phylogenetic tree based on concatenation of 26 housekeeping genes of all publicly available *Microcystis* genomes as well as metagenomes from the WLECC. Groups shown by color were previously defined and described by Dick et al., 2021. WLE strains are distributed throughout the tree, clustering with groups Mae1, Mae5, Mf1_ael, Mf1_ae3, Mae4, and Mae3. WLE strains are bolded and denoted with a green circle.

Within WLE isolates, the lowest average nucleotide identity (ANI) value between two strains was between LE19-84.1 and LE19-8.1 (95.608%) and the highest ANI was between LE17-10 and LE19-4.1 (99.958%). Hierarchical clustering based on pairwise ANI scores revealed that the WLE Culture Collection comprises two major groups. Within each group, there are several subgroups that have greater than 99% ANI, suggesting closely related or identical strains. For example, in group 2, isolates LE17-10, LE19-4.1, and LE19-196.1 share over 99.9% ANI (Fig. 2). These isolates were collected from various locations throughout the lake, during different months of the year, and in the case of LE17-10, years apart from each other (Table 1), indicating that closely related *Microcystis* strains persist through time and space in WLE. Generally, phylogenomic analyses agree with hierarchical clustering based on pairwise ANI scores (Fig 2,3). Phylogenomic analyses indicate 6-7 distinct clades of *Microcystis* isolated from WLE, further highlighting the extent of the genetic diversity within this system.

**Figure 2:**
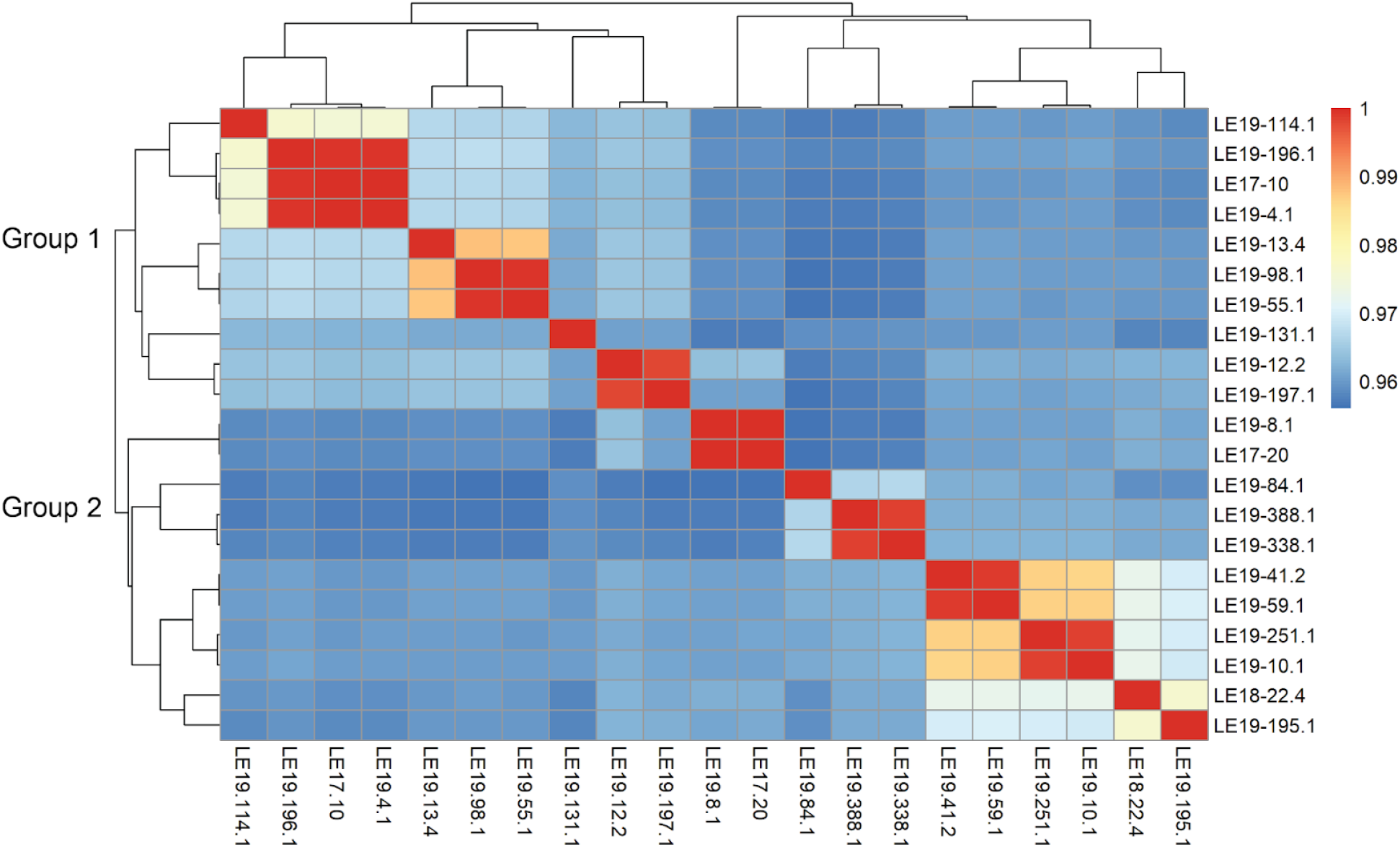
Heatmap of the Average Nucleotide Identity pairwise comparisons for the WLE *Microcystis* strains.

**Figure 3:**
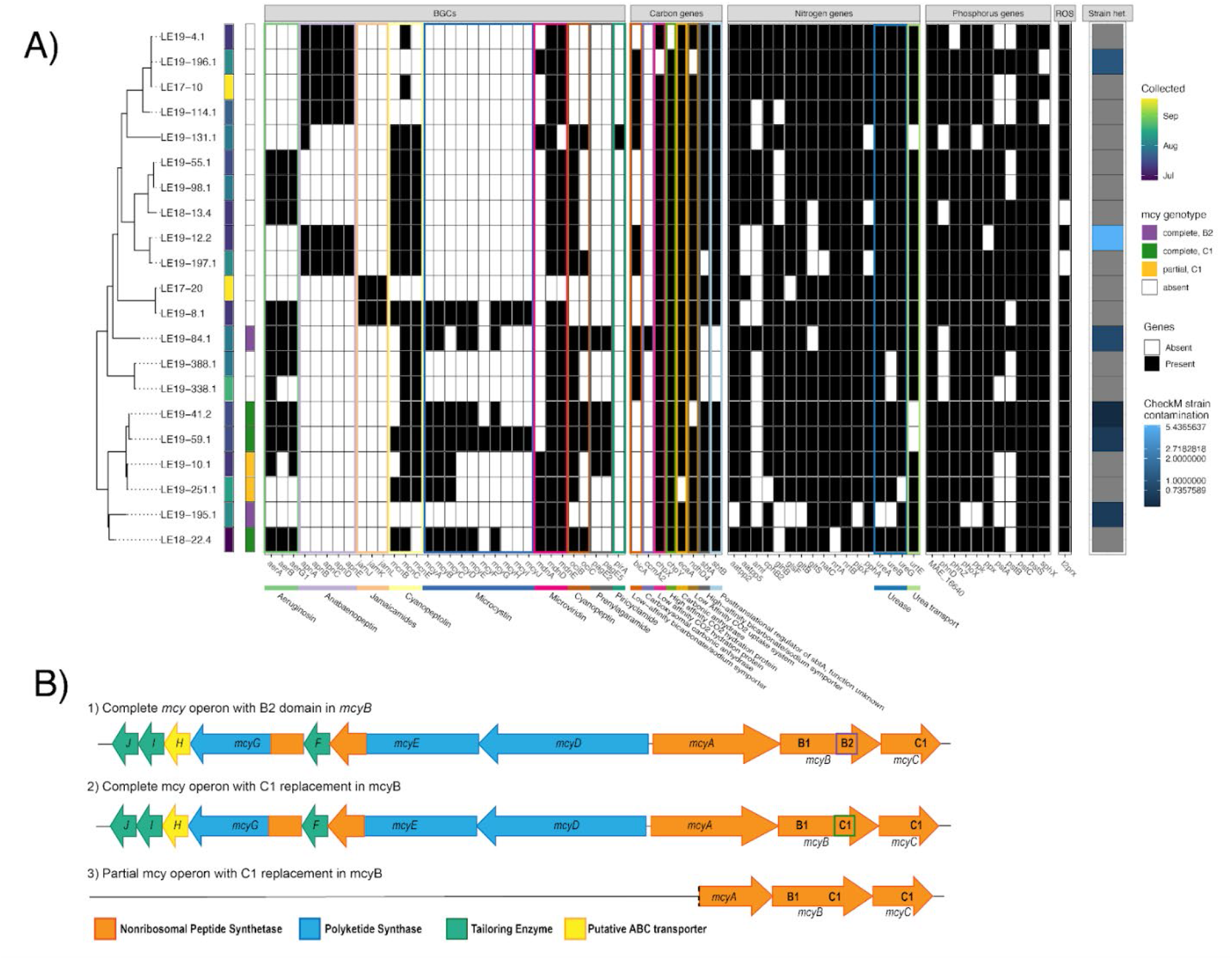
WLE *Microcystis* strains differ in gene content (A). The presence/absence of various genes is shown in relationship to strain phylogeny inferred from the concatenated housekeeping gene tree (Fig. 1). Genes that were present are colored black and absent genes are white for each culture. Strain contamination (contamination * strain heterogeneity) determined with CheckM is shown on the right, as strain heterogeneity directly reported by CheckM corresponds to the proportion of contamination predicted to originate from closely related organisms. To overcome assembly and genomic binning inefficiencies, reads from each sample were mapped to *mcy* genes to assess the true genotype present. The partial genotype (yellow) includes truncated *mcyA* and complete *mcyB* and *mcyC*, while the complete genotype (green and purple) includes all *mcy* genes. B) Gene schematics for detected *mcy* operon genotypes. B1) 2 WLE *Microcystis* strains had the complete mcy operon with the B2 domain intact in *mcyB* (purple). B2) 4 WLE *Microcystis* strains contained a complete *mcy* operon with a C1 replacement in *mcyB* (green). B3) 2 strains contained a partial *mcy* genotype in which part of *mcyA*, and complete *mcyB* and *mcyC* genes were intact. This genotype also had a C1 replacement.

We next analyzed the WLE *Microcystis* genomes for certain genes of functional interest that are related to toxin production and/or bloom progression. The majority of strains (14/21 strains) lacked *mcy* genes, and therefore microcystin producing potential. These strains formed a large clade that was monophyletic with the exception of LE19-84.1, which contains a complete *mcy* operon (Fig 3). The remaining 6 strains form another monophyletic group, with four containing the complete *mcy* operon and 2 that have a partial genotype, in which a partial *mcyA*, complete *mcyB*, and *mcyC* are present (Yancey et al., 2022a) (Fig. 3). Of the strains that had the complete *mcy* operon, 2 strains, LE19-84.1 and LE19-195.1, contained a *mcyB* gene with a B2 domain, while 3 strains with the complete *mcy* operon had a *mcyB* with a C1 replacement, which impacts the type of congener produced (Mikalsen et al., 2003). LE19-10.1 and LE19-251.1, which contained the partial genotype, both contained an *mcyB* gene with a C1 replacement (Fig. 3B).

Broader gene content is also varied across *Microcystis* isolates, with gene presence and absence generally being consistent within clades (Fig 3). There was greatest phylogenetic conservation in the presence of genes for decomposition of reactive oxygen species (ROS), nitrogen (N) and phosphorous (P) metabolism, and carbon concentrating mechanisms (CCM). None of the WLE *Microcystis* isolates contained *katG*, an enzyme used for the degradation of hydrogen peroxide (Bernroitner et al., 2009); *katG* was previously shown to be absent from most, but not all cells, in natural WLE *Microcystis* populations (Smith et al., 2022a). However, *t2prx*, a type 2 peroxiredoxin (Schuurmans et al., 2018), was present in every strain analyzed. Generally, each *Microcystis* strain contained the majority of N and P genes essential for primary metabolism and growth. However, only 6 isolates (LE17-10, LE18-22.4, LE19-196.1, LE19-4.1, LE19-55.1, and LE19-98.1) contained the ammonium transporter gene, *amt* (Fig 3). Five *Microcystis* MAGs (LE18-22.4, LE19-114.1, LE19-195.1, LE19-55.1, and LE19-98.1) are missing *glnB*, a regulator of *glnA* which encodes a glutamine synthetase (Fig 3). However, read mapping analysis identified *glnB* in all *Microcystis* genomes; this discrepancy is likely explained by failure to assemble or bin due to low read coverage (data not shown). Phosphorus uptake and metabolism genes were largely conserved across *Microcystis*, with some strains missing *pstA, pstB, pstC*, or *pstS*, which are involved in phosphate transport. However, strains that lacked one of the above listed genes contained other genes for phosphate transport (Fig 3).

We also analyzed genes encoding assimilatory pathways in the *Microcystis* genomes (Fig. 4). Many assimilatory pathways, including biosynthesis of certain amino acids (arginine, aspartate, isoleucine, leucine, valine), the Embden-Meyerhof-Parnas pathway, pentose phosphate pathway, and assimilatory ammonia incorporation and nitrate reduction, were complete across most or all strains (data not shown, Fig. 4). However, certain genes for biosynthesis of methionine were consistently absent. Genes for vitamin B12 biosynthesis were consistently present among strains but they appear to lack the capacity for biosynthesis of thiamin (B1).

**Figure 4:**
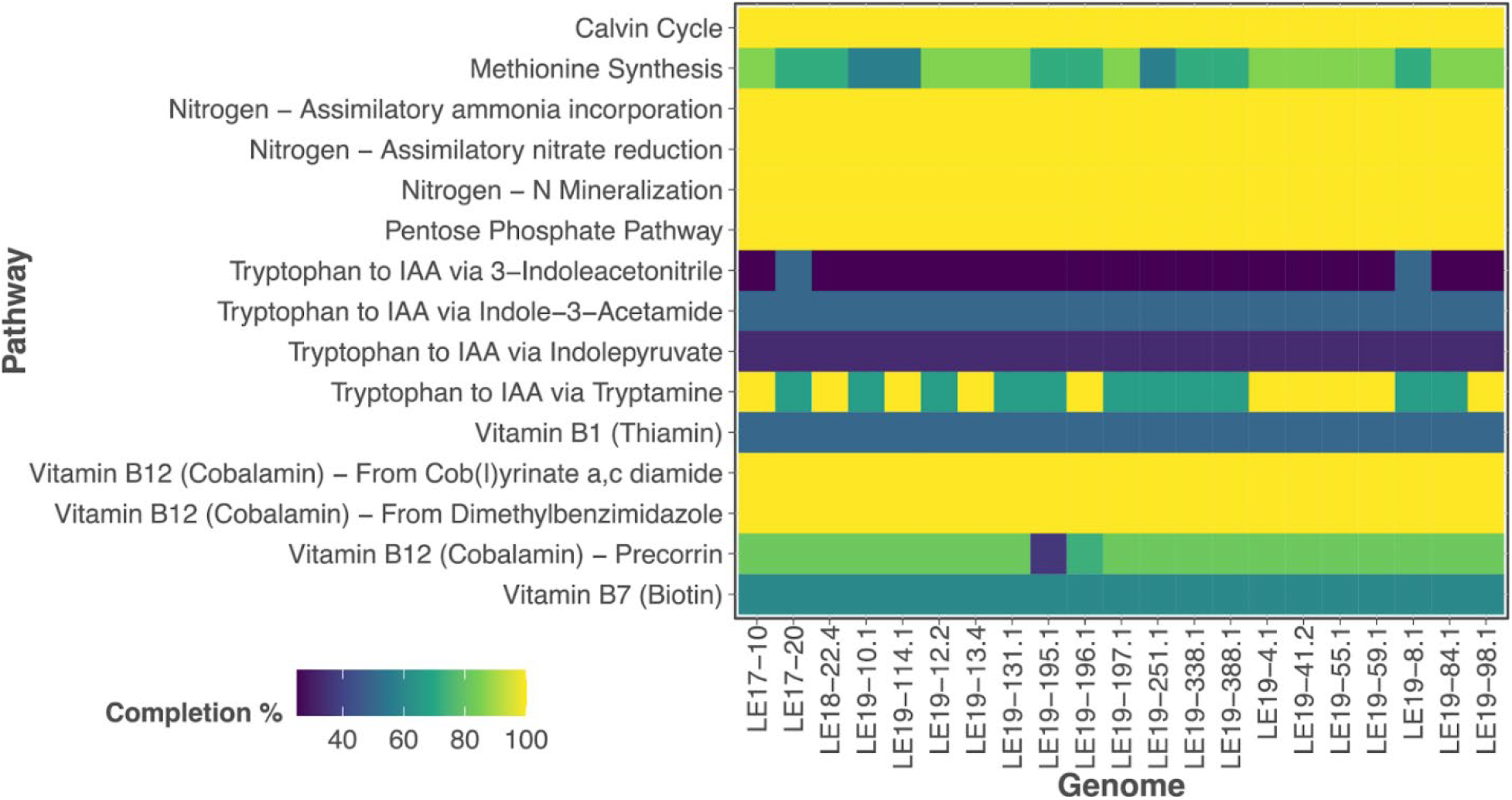
Select metabolic functional pathways for each *Microcystis* isolate. Pathway completion percentage is defined as the ratio of genes present for a pathway compared to the total genes required.

CCM gene presence and absence was also investigated as variable CCM genotypes can impact fitness in changing bloom conditions (M et al., 2005; Raven et al., 2020; Sandrini et al., 2014). The majority of CCM genes were conserved with some variability in bicarbonate uptake genes *bicA*, *sbtA*, and *sbtB*, which have been suggested to favor certain environmental conditions (Sandrini et al, 2014, Visser et al., 2016). Of the 21 strains, 5 isolated during the initiation of the cyanobacterial bloom contained *bicA*, *sbtA*, and *sbtB*, and are therefore considered generalist species able to take advantage of inorganic carbon across a wide range of concentrations. Most strains (12/21) were high bicarbonate affinity specialists, and contained only *sbtA* and *sbtB* (1 additional strain contained only *sbtB*). These strains were isolated over a wide temporal period from pre-bloom until bloom decline. Only 3 strains, LE19-197.1, LE19-388.1, and LE19-84.1, contained solely *bicA* and are considered high bicarbonate flux specialists; these were isolated during the peak of the bloom. *CcmA2*, a specialized carboxysome that catalyzes the reaction of bicarbonate to carbon dioxide (Ogawa et al., 1994), was absent from every strain except LE19-84.1. Regardless, all strains had *ccmA1*, which is an isoform of *ccmA2*, and therefore the presence of both may be functionally redundant.

Genes responsible for the synthesis of secondary metabolites were also investigated, as they can be highly variable across *Microcystis* strains and be responsible for the production of potent cyanotoxins (Dick et al., 2021; Dittmann et al., 2015; Pérez-Carrascal et al., 2019). All strains analyzed contained genes from at least one common *Microcystis* biosynthetic gene cluster (BGC). Aeruginosin and cyanopeptolin genes, which encode known cyanotoxins (Ishida et al., 2009, 1999; Martin et al., 1993; Rounge et al., 2007) were commonly detected in WLE *Microcystis* strains, observed in 12 and 16 of the 21 genomes respectively. Genes for production of microviridin B were also common–all but 3 isolates had *mdnB* and *mdnD*, while 11/21 had *mdnA*. Piricyclamide producing genes *pirA* and *pirG*, which have dynamic gene rearrangement and order (Leikoski et al 2012, Welker and Dohren 2008), were more varied among WLE strains in both gene content and sequence similarity. Variation of these genes suggests they may encode for compounds related to piricyclamide rather than the piricyclamide cluster itself. Notably, LE17-20 had no substantial hit to any BGCs screened suggesting it either has minimal biosynthetic potential, or more likely, that its genome encodes for other BGCs not analyzed in this study.

### 3.3 Abundance and Composition of Microcystins

The 5 strains that contained a complete *mcy* operon produced detectable levels of particulate MCs via ELISA measurement (Fig 5A). The 2 strains that contained partial *mcy* operons did not produce MCs above the limit of detection (0.1 μg/L) (Fig 5A). MC concentrations varied between two independent measurements, suggesting that production varies over the course of cell growth. The 14 strains lacking *mcy* genes, and therefore MC producing potential, did not contain quantifiable amounts of MC with the exception of LE19-55.1 and LE19-98.1. Detection of MC was observed for these strains in both independent measurements (Fig 5A). Read mapping revealed the presence of 10 reads that mapped to the *mcy* operon from LE19-98.1 (Table S4) indicating potential low level “contamination” of a MC producing strain within this culture. No reads from LE19-55.1 mapped to the *mcy* operon (Table S4) indicating the possibility of contamination from an unknown source.

**Figure 5:**
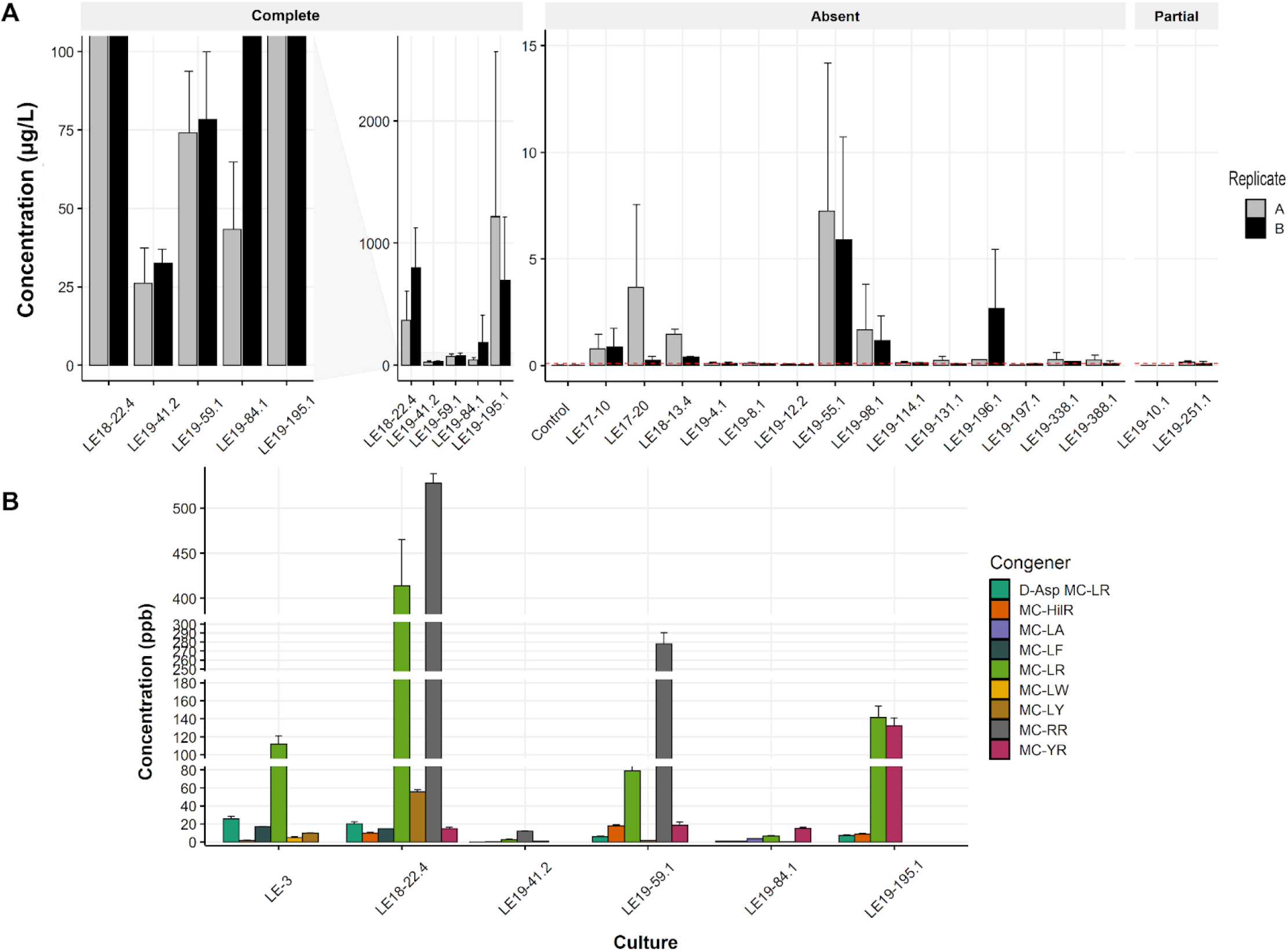
A) Measured particulate microcystin concentrations in all WLE Culture Collection isolates as measured via ELISA. Strains that produced lower amounts of MCs (<100 ug/uL) are emphasized in the left panel while all strains are depicted in the right panel for the complete genotype. Two separate replicate measurements were completed for ELISA analysis. The limit of detection, 0.1 ug/L, is shown on the absent and partial plots with a dotted red line. B) Congener detection and quantification of MCs from LE-3 and the 5 WLE culture isolates with the complete *mcy* operon.

We further analyzed the 5 WLE strains with complete *mcy* operons by LC-MS to characterize the microcystin congeners produced. *Microcystis* isolate LE-3 (Brittain et al., 2000) was included in the analysis for comparison. All 9 congeners screened for were detected in at least one strain with MC-LR being the most common and abundant followed by MC-RR and - YR. MC-LR, -RR, -LA, and -YR are the most abundant congeners observed in WLE cyanoHABs in recent years (Chaffin et al., 2021; Palagama et al., 2020). At the time of analysis, LE18-22.4 was producing the greatest amount of MC per congener (over 400 ppb for LR) as well as the greatest diversity with 7 detectable congeners. LE19-84.1 and LE19-41.2 produced 2-4 congeners at detectable levels and were much lower in their concentration compared to other strains (<40 ppb, Fig 5B). In addition to MC-LR, -RR, -LA, and -YR, MC-DAsp-LR, HilR, and - LF were detected in strain LE18-22.4. Production of these congeners in culture may be possible due to the abundance of diverse substrates in nutrient replete media and the flexibility of the substrate binding complex in MC biosynthesis enzymes (Tooming-Klunderud et al., 2008).

### 3.4 Associated Bacteria

*Non-Microcystis* bacteria within the WLE cultures were diverse, consisting of 34 genera. No two cultures contained identical bacterial communities (Fig 6) even though some *Microcystis* strains are nearly genetically identical (Fig 2). The most common associated heterotrophic bacteria identified across the cultures were known MC degrader *Novosphingobium* sp. (10/22 cultures), (Dziga et al., 2013), *Rhabdaerophilium* sp. (5/22 cultures), and *Silanimonas* sp. (4/22 cultures) as determined by mapping reads to MAGs obtained from each sample (Fig 6). However, taxonomic annotation of sample reads with the k-mer profiling tools Kraken and Bracken (Lu et al., 2017; Wood and Salzberg, 2014) found that *Erythobacter* (17/22), *Novosphingobium* (12/22), and *Flavobacterium* (12/22) were most abundant across cultures (Fig. S2). In many cases, the associated bacteria were more abundant than the *Microcystis* strains present within each culture (Fig 5, Table S1), though biases in DNA extraction could influence this result. Associated bacterial genomes were also queried for catalase genes, which degrade hydrogen peroxide, thus ameliorating ROS stress on *Microcystis* (Morris et al., 2008; Smith et al., 2022a). Of the 22 cultures, 17 cultures contained organisms with catalase genes.

**Figure 6:**
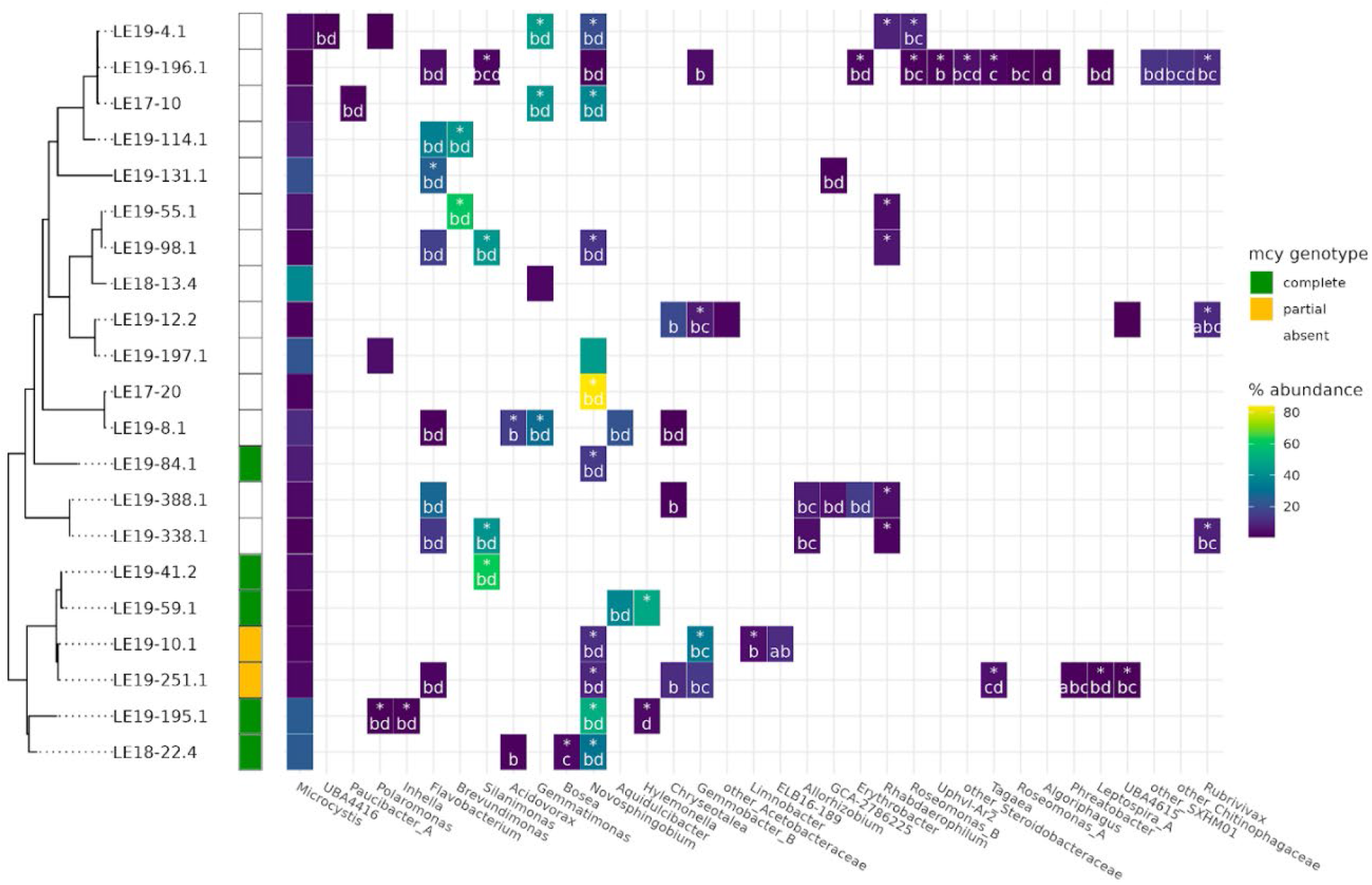
Taxonomy, relative abundance, and key functions of bacteria for which metagenome assembled genomes were retrieved in each *Microcystis* culture. Some cultures only contain 1 heterotroph (LE17-20) while others contain more than 10 genera (LE19-196.1). Asterisks indicate the presence of catalase genes and a-d represents the presence of respective *mlr* genes, part of a microcystin degradation pathway.

We next examined the genomic potential of associated bacteria to degrade cyanotoxins. Nineteen cultures were shown to contain *mlr* genes, which represent a known mechanism to degrade MCs (Bourne et al., 2001), but no MAGs contained complete *mlr* operons (*mlrA-D*). All four *mlr* genes were detected across multiple bins in only two mixed cultures, LE19-251.1 and LE19-10.1 which respectively have partial and absent *mcy* genotypes (Fig 6). Both *mlrB* and *mlrC* degrade the linearized microcystin product (Dziga et al., 2013) and *mlrC* is capable of linearizing MCs similar to *mlrA* (Dziga et al., 2012; Shimizu et al., 2012), suggesting that these bacteria may use intermediate breakdown products as substrates. Of the 7 *Microcystis* strains containing complete or partial sets of *mcy* genes, *mlr* genes were detected in all cultures. *MlrD*, a putative microcystin transporter or membrane anchoring protein (Bourne et al., 2001), was present in 18 cultures, including all that contained *mcy* genes (Fig 6). The associated bacteria that contained *mlr* genes were distinct across cultures, suggesting multiple taxa have the ability to degrade MC. *MlrA* was detected in three cultures, two with a partial *mcy* operon–LE19-251.1 (*Phreatobacter*), LE19-10.1 (*ELB16-189*), and LE19-12.2 (*Rubrivivax). MlrC*, which can also linearize cyclic MCs (Dziga et al., 2012; Shimizu et al., 2012), was detected in 8 cultures, including both *Microcystis* strains with the partial *mcy* operon and one *Microcystis* strain (LE18-22.4) with the complete *mcy* operon (Fig 6). The most common associated bacteria identified with *mlr* genes were *Novosphingobium* (11 cultures with *mlrB* and *mlrD*)), *Flavobacterium* (8 cultures with *mlrB* and *mlrD*), Chryseotalea (4 cultures with *mlrB* and one with *mlrB and mlrD*), *Gemmobacter* B (3 cultures with *mlrB* and *mlrC* and one with *mlrB*), and *Silanimonas* (3 cultures with *mlrB* and *mlrD* and one with *mlrB*-*D*).

## 4. Discussion

Achieving axenic *Microcystis* isolates from environmental samples can be labor intensive, and also fails to capture the full diversity of organisms observed in natural communities (Schloss and Handelsman, 2005). Due to extensive genetic diversity observed between *Microcystis* strains (Dick et al., 2021; Harke et al., 2016; Humbert et al., 2013), the choice of axenic *Microcystis* culture for experimentation is critical. Special care should be taken when selecting axenic *Microcystis* isolates from culture collections as these strains may not represent common genotypes associated with certain bodies of water. Here, we report the isolation, initial characterization, and genome sequencing of 21 strains of *Microcystis* from WLE, comprising the WLECC. Although isolation and cultivation methods were largely the same from year to year, we isolated a diverse collection of *Microcystis* strains with variable accessory genomes and associated bacterial communities that represent the extensive diversity of *Microcystis* strains (Dick et al., 2021) and colony-associated bacteria (Smith et al. 2021) observed in the field. Whereas the 159 publicly available genomes of cultured *Microcystis* are dominated by genotypes with the complete set of *mcy* genes (Dick et al. 2021), with only one partial genotype (Yancey et al. 2022), the majority of WLECC strains lack *mcy* genes altogether, and two strains contain the partial genotype. Further, the WLECC includes a diverse array *Microcystis* traits (secondary metabolism, CCM, nutrient metabolism) that may be critical in strain succession and community function in natural environments (Dick et al., 2021). Thus, this collection is a valuable resource for studies of *Microcystis* strain diversity and microbial dynamics in WLE and for the greater cyanoHAB research community as it provides an intermediate between complex environmental samples and pure axenic culture.

Although WLECC cultures contain simplified bacterial consortia relative to complex environmental samples, we encountered challenges in metagenomic data analysis, resulting in draft genome sequences of varying quality (Table S3). However, the majority of *Microcystis* MAGs generated for this study are high quality, with longer contiguous sequence (Table S3) compared to MAGs from environmental samples (Yancey et al., 2022b). Lower-quality draft genomes provide valuable insight into gene content and microbial community diversity, but deeper sequencing with long-read technologies would improve assembly quality and enable definitive conclusions about gene absence, gene order, and overall genome architecture. Additionally, genome sequencing revealed that some isolates determined to be “unialgal” via 16S rRNA gene sequencing actually contained genetic diversity at a subspecies level. This was evident by the low abundance of *mcy* operon reads (<10, Table S2) in genomes reported to have the absent genotype (Fig 3), detectable levels of MCs in those isolates (Fig 5A), and detectable strain heterogeneity via CheckM (Fig 3, Table S3).

The genome sequencing results in this study underscore the substantial genetic diversity observed in *Microcystis* at a subspecies level, highlighting our limited understanding of the extent of genotypic and ecophysiological diversity of this genus. The 21 isolates in the WLECC represent 12 unique strains of *Microcystis* (Fig 2,3) based on a cut off of 99.8% ANI and phylogenetic clustering. Of these 12 proposed strain groups, 6 are represented by more than one isolate and 6 are singletons (Table 1). The WLECC isolates are scattered phylogenetically across the *Microcystis* tree, interspersed with *Microcystis* isolated from around the world (Fig 1), consistent with global dispersal of *Microcystis* strains (Marmen et al., 2016; Moreira et al., 2014). Taken together with our finding of near-identical strains at different stations and years, these results are consistent with the notion of a limited pool of *Microcystis* strains with wide spatial and temporal distribution. The WLECC strains are genetically diverse, with variability in N and P metabolism, CCMs, and secondary metabolite genes (Fig 3) suggesting each strain is specifically adapted to micro-niches resulting from spatiotemporal environmental gradients observed in WLE (Cooperative Institute for Great Lakes Research, n.d.; Dick et al., 2021).

The sequencing results also reveal our limited understanding of the multi-faceted controls on differential MC congener production, and the need for further studies to constrain the role of both genotype and associated bacteria. *Microcystis* has the ability to produce a wide range of MC congeners with varying degrees of toxicity (Chernoff et al., 2020; Bouaïcha et al., 2019; Díez-Quijada et al., 2019). Our findings are consistent with previous work that demonstrates the ability of *mcyB* B2 genotypes to produce at least MC-LR, while C1 genotypes may produce at least MC-LR, and MC-RR (Fig. 5B) (Mikalsen et al., 2003). Our results also reveal the ability of these strains to produce several other congeners including MC-LA and MC-YR, which are observed in natural WLE cyanoHABs along with -LR and -RR (Chaffin et al., 2021, Palagama et al., 2020), as well as others not commonly observed (-LF, -HilR) (Fig 5B). Previous studies have established *mcy* operon architecture as an important component of differential congener production (Mikalsen et al., 2003; Tooming-Klunderud et al., 2008). However, several other factors can influence congener production such as flexibility of the substrate binding complex (Tooming-Klunderud et al., 2008), nitrogen form and availability (Puddick et al., 2014), and available intracellular amino acid substrates (Tonk et al., 2008). Since measurements were completed in culture, it is unlikely that these abiotic variables, which were controlled for, have the greatest effect on differential congener production between closely related strains. Instead, different assemblages of associated bacteria may be responsible for observed differences in congener production. For example, LE19-195.1 and LE18-22.4 are genetically similar (over 99.8% ANI) yet produce varying ratios of the 9 measured MC congeners (Fig 5B). While it is possible these differences may be attributed to undiscovered *mcy* genotypes, these cultures also contain distinct consortia of associated bacteria (Fig. 6) which may impact substrate availability and metabolic exchange (Baran et al., 2015).

Indeed, the gene content of the WLE *Microcystis* strains suggests possible important interactions with bacteria. For example, the lack of genes for methionine biosynthesis may indicate auxotrophy, and previous work has implicated heterotrophic bacteria in supplying *Microcystis* with methionine (Jung et al., 2022). In turn, the conserved capacity for vitamin B12 biosynthesis across all strains is consistent with evidence that vitamin B12 from *Microcystis* may support heterotrophic bacteria (Smith et al., 2022b). The lack of genes for vitamin B1 (thiamin) biosynthesis in WLE *Microcystis* strains parallels findings in HAB-forming dinoflagellate species, which are suggested to be thiamin auxotrophs (Tang et al., 2010). Cyanobacterial blooms have been implicated in interfering with trophic transfer of B1 (Fridolfsson et al., 2018), but we are not aware of previous studies showing that *Microcystis* lacks B1 biosynthesis.

The WLECC provides a unique means to further investigate this and other possible mutualistic interactions between *Microcystis* and their microbiome. The WLECC captures a breadth of diversity of physically associated bacteria, with minimal overlap between cultures, even for *Microcystis* strains that are genetically similar (Fig 6). This agrees with previous work that showed WLE *Microcystis* colonies have different associated bacteria between strains and across time, lacking a “core” bacterial community found across all strains (Smith et al., 2021). While *katG*, an enzyme used by cyanobacteria to degrade hydrogen peroxide (Bernroitner et al., 2009; Mutsuda et al., 1996), was undetected in all 21 *Microcystis* strains, a large portion of associated bacteria in these cultures contained catalase genes, suggesting they are likely important in alleviating ROS stress from the cyanoHAB phycosphere. During periods of rapid growth in WLE cyanoHABs, concentrations of hydrogen peroxide can reach high levels of 600-800 nM (Cory et al., 2016; Smith et al., 2022a), which can cause oxidative stress and cell death in microorganisms (Imlay, 2003; Latifi et al., 2009). Our results are consistent with previous studies suggesting “helper” heterotrophic bacteria are critical in facilitating rapid growth of cyanobacteria in aquatic systems (Morris et al., 2008; Smith et al., 2022a) by relieving ROS stress.

In contrast to universal presence of catalase genes among associated heterotrophs, the distribution of *mlr* genes for MC degradation was patchy with no bacterial genome or culture containing a complete suite of *mlr* genes (Fig. 6). Further, there was no obvious relationship between *mcy* genotype and *mlr* genes present (Fig. 6). Previous studies have shown actinobacteria (Manage et al., 2009) and a variety of bacteria isolated from WLE (Krishnan et al., 2020) are able to degrade MCs despite lacking *mlr* genes, suggesting undescribed alternative pathways that can biodegrade MCs (Dziga et al., 2017). Additionally, most cultures lack *mcy* genes and MC producing potential, but still encode for the synthesis of other secondary metabolites (Fig 3). A recent study has demonstrated the ability of *mlr*-bacteria to degrade both MCs and other cyanobacterial peptides (Santos et al., 2021) suggesting that associated bacteria within the WLECC may be capable of degrading other cyanopeptides beside MC through unknown pathways. Future research should address the substrate specificity of *mlr-*encoded enzymes, as well as cryptic pathways beyond *mlr* that can degrade MCs and/or other cyanobacterial peptides.

## 5. Conclusion

The WLECC will be a valuable resource for several directions of cyanoHAB research. First, the diversity and variable gene content of strains present opportunities to study how various physiological traits are distributed across phylogenetic clusters of *Microcystis* and how these traits are underpinned at the genetic level. Second, cultures of *Microcystis* strains with variable arrays of biosynthetic gene cluster content enable studies of the toxicology of diverse secondary metabolites from *Microcystis* (Janssen, 2019) and efforts to link “orphan” BGCs (Humbert et al., 2013; Yancey et al., 2022b) with secondary metabolite products. The cultures may also be used to study potential toxicological synergisms of various combinations of secondary metabolites (Fernandes et al., 2019; Kust et al., 2020; Pawlik-Skowrońska and Bownik, 2022), which often occur in complex mixtures in cyanobacterial blooms (Beversdorf et al., 2018, 2017). Third, this collection can be used to address how different assemblages of associated bacteria can impact aspects of growth, toxin production and degradation, and colony formation for genetically similar/identical strains of *Microcystis*. For example, LE17-10, LE19-196.1, and LE19-4.1 all belong to strain group 1 (Table 1), but these cultures consist of very different combinations of heterotrophic bacteria, with only *Novosphingobium* sp. being present in all three. Finally, the WLECC consists of strains isolated from the Great Lakes, which has previously been represented solely by LE-3. Thus, we expect the WLECC to serve as a valuable resource for bridging field-based observations with laboratory-controlled experiments to characterize the dynamics of diverse *Microcystis* strains and their microbiomes.

## Acknowledgements

We would like to thank Robert Hein for bioinformatic support. We also thank the field crew at CIGLR/GLERL including Paul Den Uyl, Dack Stuart, Kent Baker, and Holly Kelchner for allowing us to sample with them and assisting in field measurements. We would like to thank H. Kelchner and D. Gossiaux for assistance with ELISA assays. This work was supported by the Cooperative Institute of Great Lake Research (NA17OAR4320152), Air Force Office of Scientific Research grant (DURIP 14RT0605), and Lawrence Livermore National Laboratory LDRD project 20-ERD-061. Part of this work was carried out at LLNL under Contract DE-AC52-07NA27344.

## References

Allen, M.M., 1968. Simple Conditions for Growth of Unicellular Blue-Green Algae on Plates1, 2. J. Phycol. 4, 1–4. https://doi.org/10.1111/j.1529-8817.1968.tb04667.x

Alneberg, J., Bjarnason, B.S., de Bruijn, I., Schirmer, M., Quick, J., Ijaz, U.Z., Loman, N.J., Andersson, A. F., Quince, C., 2013. CONCOCT: Clustering cONtigs on COverage and ComposiTion.

Baliu-Rodriguez, D., Peraino, N.J., Premathilaka, S.H., Birbeck, J.A., Baliu-Rodriguez, T., Westrick, J.A., Isailovic, D., 2022. Identification of Novel Microcystins Using High-Resolution MS and MSn with Python Code. Environ. Sci. Technol. 56, 1652–1663. https://doi.org/10.1021/acs.est.1c04296

Baran, R., Brodie, E.L., Mayberry-Lewis, J., Hummel, E., Da Rocha, U.N., Chakraborty, R., Bowen, B. P., Karaoz, U., Cadillo-Quiroz, H., Garcia-Pichel, F., Northen, T.R., 2015. Exometabolite niche partitioning among sympatric soil bacteria. Nat. Commun. 2015 61 6, 1–9. https://doi.org/10.1038/ncomms9289

Bates, S.S., Douglas, D.J., Doucette, G.J., Léger, C., 1995. Enhancement of domoic acid production by reintroducing bacteria to axenic cultures of the diatom Pseudo-nitzschia multiseries. Nat. Toxins 3, 428–435. https://doi.org/10.1002/nt.2620030605

Bernroitner, M., Zamocky, M., Furtmüller, P.G., Peschek, G.A., Obinger, C., 2009. Occurrence, phylogeny, structure, and function of catalases and peroxidases in cyanobacteria. J. Exp. Bot. 60, 423–440. https://doi.org/10.1093/jxb/ern309

Berry, M.A., Davis, T.W., Cory, R.M., Duhaime, M.B., Johengen, T.H., Kling, G.W., Marino, J.A., Den Uyl, P.A., Gossiaux, D., Dick, G.J., Denef, V.J., 2017. Cyanobacterial harmful algal blooms are a biological disturbance to Western Lake Erie bacterial communities. Environ. Microbiol. 19, 1149–1162. https://doi.org/10.1111/1462-2920.13640

Beversdorf, L.J., Rude, K., Weirich, C.A., Bartlett, S.L., Seaman, M., Kozik, C., Biese, P., Gosz, T., Suha, M., Stempa, C., Shaw, C., Hedman, C., Piatt, J.J., Miller, T.R., 2018. Analysis of cyanobacterial metabolites in surface and raw drinking waters reveals more than microcystin. Water Res. 140, 280–290. https://doi.org/10.1016/j.watres.2018.04.032

Beversdorf, L.J., Weirich, C.A., Bartlett, S.L., Miller, T.R., 2017. Variable Cyanobacterial Toxin and Metabolite Profiles across Six Eutrophic Lakes of Differing Physiochemical Characteristics. Toxins 9, 62. https://doi.org/10.3390/toxins9020062

Binding, C.E., Stumpf, R.P., Shuchman, R.A., Sayers, M.J., 2020. Advances in Remote Sensing of Great Lakes Algal Blooms, in: Crossman, J., Weisener, C. (Eds.), Contaminants of the Great Lakes, The Handbook of Environmental Chemistry. Springer International Publishing, Cham, pp. 217–232. https://doi.org/10.1007/698_2020_589

Bingham, M., Sinha, S., Lup, F., 2015. Economic Benefits of Reducing Harmful Algal Blooms in Lake Erie.

Bouaïcha, N., Miles, C.O., Beach, D.G., Labidi, Z., Djabri, A., Benayache, N.Y., Nguyen-Quang, T., 2019. Structural diversity, characterization and toxicology of microcystins. Toxins 11, 714. https://doi.org/10.3390/toxins11120714

Bourne, D.G., Riddles, P., Jones, G.J., Smith, W., Blakeley, R.L., 2001. Characterisation of a gene cluster involved in bacterial degradation of the cyanobacterial toxin microcystin LR. Environ. Toxicol. 16, 523–534. https://doi.org/10.1002/tox.10013

Breitwieser, F.P., Baker, D.N., Salzberg, S.L., 2018. KrakenUniq: confident and fast metagenomics classification using unique k-mer counts. Genome Biol 19, 198. https://doi.org/10.1186/s13059-018-1568-0

Bridgeman, T.B., Chaffin, J.D., Filbrun, J.E., 2013. A novel method for tracking western Lake Erie Microcystis blooms, 2002–2011. J. Gt. Lakes Res. 39, 83–89. https://doi.org/10.1016/J.JGLR.2012.11004

Brittain, S.M., Wang, J., Babcock-Jackson, L., Carmichael, W.W., Rinehart, K.L., Culver, D.A., 2000. Isolation and Characterization of Microcystins, Cyclic Heptapeptide Hepatotoxins from a Lake Erie Strain of Microcystis aeruginosa. J. Gt. Lakes Res. 26, 241–249. https://doi.org/10.1016/S0380-1330(00)70690-3

Bushnell, B., n.d. BBTools User Guide - DOE Joint Genome Institute [WWW Document]. URL https://sourceforge.net/projects/bbmap/ (accessed 4.9.21).

Cao, H., Xu, D., Zhang, T., Ren, Q., Xiang, L., Ning, C., Zhang, Y., Gao, R., 2022. Comprehensive and functional analyses reveal the genomic diversity and potential toxicity of Microcystis. Harmful Algae 113, 102186. https://doi.org/10.1016/j.hal.2022.102186

Chaffin, J.D., Bratton, J.F., Verhamme, E.M., Bair, H.B., Beecher, A.A., Binding, C.E., Birbeck, J.A., Bridgeman, T.B., Chang, X., Crossman, J., Currie, W.J.S., Davis, T.W., Dick, G.J., Drouillard, K.G., Errera, R.M., Frenken, T., MacIsaac, H.J., McClure, A., McKay, R.M., Reitz, L.A., Domingo, J.W.S., Stanislawczyk, K., Stumpf, R.P., Swan, Z.D., Snyder, B.K., Westrick, J.A., Xue, P., Yancey, C.E., Zastepa, A., Zhou, X., 2021. The Lake Erie HABs Grab: A binational collaboration to characterize the western basin cyanobacterial harmful algal blooms at an unprecedented high-resolution spatial scale. Harmful Algae 108, 102080. https://doi.org/10.1016/J.HAL.2021.102080

Chaumeil, P.-A., Mussig, A.J., Hugenholtz, P., Parks, D.H., 2019. GTDB-Tk: a toolkit to classify genomes with the Genome Taxonomy Database. Bioinformatics. https://doi.org/10.1093/bioinformatics/btz848

Cooperative Institute for Great Lakes Research, U. of M.N.G.L.E.R.L., n.d. Physical, chemical, and biological water quality monitoring data to support detection of Harmful Algal Blooms (HABs) in western Lake Erie, collected by the Great Lakes Environmental Research Laboratory and the Cooperative Institute for Great Lakes Research since 2012 [WWW Document]. https://doi.org/10.25921/11da-3x54

Cory, R.M., Davis, T.W., Dick, G.J., Johengen, T., Denef, V.J., Berry, M.A., Page, S.E., Watson, S.B., Yuhas, K., Kling, G.W., 2016. Seasonal Dynamics in Dissolved Organic Matter, Hydrogen Peroxide, and Cyanobacterial Blooms in Lake Erie. Front. Mar. Sci. 3.

Den Uyl, P.A., Thompson, L.R., Errera, R.M., Birch, J.M., Preston, C.M., Ussler, W., Yancey, C.E., Chaganti, S.R., Ruberg, S.A., Doucette, G.J., Dick, G.J., Scholin, C.A., Goodwin, K.D., 2022. Lake Erie field trials to advance autonomous monitoring of cyanobacterial harmful algal blooms. Front. Mar. Sci. 9.

Dick, G.J., 2018. Genomic Approaches in Earth and Environmental Sciences. John Wiley & Sons, Ltd. https://doi.org/10.1002/9781118708231

Dick, G.J., Duhaime, M.B., Evans, J.T., Errera, R.M., Godwin, C.M., Kharbush, J.J., Nitschky, H.S., Powers, M.A., Vanderploeg, H.A., Schmidt, K.C., Smith, D.J., Yancey, C.E., Zwiers, C.C., Denef, V.J., 2021. The genetic and ecophysiological diversity of *Microcystis*. Environ. Microbiol. 1462-2920.15615. https://doi.org/10.1111/1462-2920.15615

Díez-Quijada, L., Prieto, A.I., Guzmán-Guillén, R., Jos, A., Cameán, A.M., 2019. Occurrence and toxicity of microcystin congeners other than MC-LR and MC-RR: A review. Food Chem. Toxicol. 125, 106–132. https://doi.org/10.1016/J.FCT.2018.12.042

Dittmann, E., Gugger, M., Sivonen, K., Fewer, D.P., 2015. Natural Product Biosynthetic Diversity and Comparative Genomics of the Cyanobacteria. Trends Microbiol. 23, 642–652. https://doi.org/10.1016/J.TIM.2015.07.008

Dziga, D., Maksylewicz, A., Maroszek, M., Budzyńska, A., Napiorkowska-Krzebietke, A., Toporowska, M., Grabowska, M., Kozak, A., Rosińska, J., Meriluoto, J., 2017. The biodegradation of microcystins in temperate freshwater bodies with previous cyanobacterial history. Ecotoxicol. Environ. Saf. 145, 420–430. https://doi.org/10.1016/j.ecoenv.2017.07.046

Dziga, D., Wasylewski, M., Szetela, A., Bocheńska, O., Wladyka, B., 2012. Verification of the Role of MlrC in Microcystin Biodegradation by Studies Using a Heterologously Expressed Enzyme. Chem. Res. Toxicol. 25, 1192–1194. https://doi.org/10.1021/tx300174e

Dziga, D., Wasylewski, M., Wladyka, B., Nybom, S., Meriluoto, J., 2013. Microbial degradation of microcystins. Chem. Res. Toxicol. 26, 841–852. https://doi.org/10.1021/tx4000045

Eddy, S.R., 2011. Accelerated Profile HMM Searches. PLoS Comput Biol 7, e1002195. https://doi.org/10.1371/journal.pcbi.1002195

Edgar, R.C., 2004. MUSCLE: multiple sequence alignment with high accuracy and high throughput. Nucleic Acids Res. 32, 1792–1797. https://doi.org/10.1093/NAR/GKH340

Eren, A.M., Esen, Ö.C., Quince, C., Vineis, J.H., Morrison, H.G., Sogin, M.L., Delmont, T.O., 2015. Anvi’o: an advanced analysis and visualization platform for ‘omics data. PeerJ 3, e1319. https://doi.org/10.7717/peerj.1319

Eren, A.M., Kiefl, E., Shaiber, A., Veseli, I., Miller, S.E., Schechter, M.S., Fink, I., Pan, J.N., Yousef, M., Fogarty, E.C., Trigodet, F., Watson, A.R., Esen, Ö.C., Moore, R.M., Clayssen, Q., Lee, M.D., Kivenson, V., Graham, E.D., Merrill, B.D., Karkman, A., Blankenberg, D., Eppley, J.M., Sjödin, A., Scott, J.J., Vázquez-Campos, X., McKay, L.J., McDaniel, E.A., Stevens, S.L.R., Anderson, R.E., Fuessel, J., Fernandez-Guerra, A., Maignien, L., Delmont, T.O., Willis, A.D., 2021. Community-led, integrated, reproducible multi-omics with anvi’o. Nat. Microbiol. 6, 3–6. https://doi.org/10.1038/s41564-020-00834-3

Fernandes, K., Gomes, A., Calado, L., Yasui, G., Assis, D., Henry, T., Fonseca, A., Pinto, E., 2019. Toxicity of Cyanopeptides from Two Microcystis Strains on Larval Development of Astyanax altiparanae. Toxins 11, 220. https://doi.org/10.3390/toxins11040220

Fridolfsson, E., Lindehoff, E., Legrand, C., Hylander, S., 2018. Thiamin (vitamin B1) content in phytoplankton and zooplankton in the presence of filamentous cyanobacteria. Limnol. Oceanogr. 63, 2423–2435. https://doi.org/10.1002/lno.10949

Griffith, A.W., Gobler, C.J., 2020. Harmful algal blooms: A climate change co-stressor in marine and freshwater ecosystems. Harmful Algae 91, 101590. https://doi.org/10.1016/j.hal.2019.03.008

Guillard, R.R.L., Lorenzen, C.J., 1972. Yellow-Green Algae with Chlorophyllide C1,2. J. Phycol. 8, 10–14. https://doi.org/10.1111/j.1529-8817.1972.tb03995.x

Harke, M.J., Steffen, M.M., Otten, T.G., Wilhelm, S.W., Wood, S.A., Paerl, H.W., 2016. A review of the global ecology, genomics, and biogeography of the toxic cyanobacterium, Microcystis spp. Harmful Algae 54, 4–20. https://doi.org/10.1016/J.HAL.2015.12.007

Hellweger, F.L., Martin, R.M., Eigemann, F., Smith, D.J., Dick, G.J., Wilhelm, S.W., 2022. Models predict planned phosphorus load reduction will make Lake Erie more toxic. Science 376, 1001–1005. https://doi.org/10.1126/SCIENCE.ABM6791

Ho, J.C., Michalak, A.M., Pahlevan, N., 2019. Widespread global increase in intense lake phytoplankton blooms since the 1980s. Nature 574, 667–670. https://doi.org/10.1038/s41586-019-1648-7

Hoke, A.K., Reynoso, G., Smith, M.R., Gardner, M.I., Lockwood, D.J., Gilbert, N.E., Wilhelm, S.W., Becker, I.R., Brennan, G.J., Crider, K.E., Farnan, S.R., Mendoza, V., Poole, A.C., Zimmerman, Z.P., Utz, L.K., Wurch, L.L., Steffen, M.M., 2021. Genomic signatures of Lake Erie bacteria suggest interaction in the Microcystis phycosphere. PLOS ONE 16, e0257017. https://doi.org/10.1371/JOURNAL.PONE.0257017

Huisman, J., Codd, G.A., Paerl, H.W., Ibelings, B.W., H Verspagen, J.M., Visser, P.M., 2018. Cyanobacterial blooms. Nat. Rev. Microbiol. https://doi.org/10.1038/s41579-018-0040-1

Humbert, J.-F., Barbe, V., Latifi, A., Gugger, M., Calteau, A., Coursin, T., Lajus, A., Castelli, V., Oztas, S., Samson, G., Longin, C., Medigue, C., de Marsac, N.T., 2013. A Tribute to Disorder in the Genome of the Bloom-Forming Freshwater Cyanobacterium Microcystis aeruginosa. PLoS ONE 8, e70747. https://doi.org/10.1371/journal.pone.0070747

Imlay, J.A., 2003. Pathways of oxidative damge. Annu. Rev. Microbiol. 57, 395–418.

Ishida, K., Okita, Y., Matsuda, H., Okino, T., Murakami, M., 1999. Aeruginosins, protease inhibitors from the cyanobacterium Microcystis aeruginosa. Tetrahedron 55, 10971–10988. https://doi.org/10.1016/S0040-4020(99)00621-3

Ishida, K., Welker, M., Christiansen, G., Cadel-Six, S., Bouchier, C., Dittmann, E., Hertweck, C., De Marsac, N.T., 2009. Plasticity and evolution of aeruginosin biosynthesis in cyanobacteria. Appl. Environ. Microbiol. 75, 2017–2026. https://doi.org/10.1128/AEM.02258-08

Janssen, E.M.L., 2019. Cyanobacterial peptides beyond microcystins – A review on co-occurrence, toxicity, and challenges for risk assessment. Water Res. 151, 488–499. https://doi.org/10.1016/j.watres.2018.12.048

Jung, J., Baek, J.H., Lee, Y., Jeong, S.E., Jeon, C.O., 2022. The Self-Bleaching Process of Microcystis aeruginosa is Delayed by a Symbiotic Bacterium Pseudomonas sp. MAE1-K and Promoted by Methionine Deficiency. Microbiol. Spectr. 10, eO181422. https://doi.org/10.1128/spectrum.01814-22

Krishnan, A., Zhang, Y., Balaban, M., Seo, Y., Mou, X., 2020. Taxonomic and Genotypical Heterogeneity of Microcystin degrading Bacterioplankton in Western Lake Erie. Harmful Algae 98, 101895. https://doi.org/10.1016/j.hal.2020.101895

Kust, A., Řeháková, K., Vrba, J., Maicher, V., Mareš, J., Hrouzek, P., Chiriac, M.-C., Benedová, Z., Tesařová, B., Saurav, K., 2020. Insight into Unprecedented Diversity of Cyanopeptides in Eutrophic Ponds Using an MS/MS Networking Approach. Toxins 12, 561. https://doi.org/10.3390/toxins12090561

Latifi, A., Ruiz, M., Zhang, C.-C., 2009. Oxidative stress in cyanobacteria. FEMS Microbiol. Rev. 33, 258–278. https://doi.org/10.1111/j.1574-6976.2008.00134.x

Li, D., Liu, C.-M., Luo, R., Sadakane, K., Lam, T.-W., 2015. MEGAHIT: an ultra-fast single-node solution for large and complex metagenomics assembly via succinct de Bruijn graph. Bioinformatics 31, 1674–1676. https://doi.org/10.1093/bioinformatics/btv033

Li, H., 2018. Minimap2: pairwise alignment for nucleotide sequences. Bioinformatics 34, 3094–3100. https://doi.org/10.1093/bioinformatics/bty191

Lu, J., Breitwieser, F.P., Thielen, P., Salzberg, S.L., 2017. Bracken: estimating species abundance in metagenomics data. PeerJ Comput Sci 3, e104. https://doi.org/10.7717/peerj-cs.104

M, G., J, B., Ja, R., 2005. CO2 concentrating mechanisms in algae: mechanisms, environmental modulation, and evolution. Annu. Rev. Plant Biol. 56. https://doi.org/10.1146/annurev.arplant.56.032604.144052

Madden, T., 2013. The BLAST Sequence Analysis Tool.

Manage, P.M., Edwards, C., Singh, B.K., Lawton, L.A., 2009. Isolation and Identification of Novel Microcystin-Degrading Bacteria. Appl. Environ. Microbiol. 75, 6924–6928. https://doi.org/10.1128/AEM.01928-09

Marmen, S., Aharonovich, D., Grossowicz, M., Blank, L., Yacobi, Y.Z., Sher, D.J., 2016. Distribution and Habitat Specificity of Potentially-Toxic Microcystis across Climate, Land, and Water Use Gradients. Front. Microbiol. 7.

Martin, C., Oberer, L., Ino, T., König, W.A., Busch, M., Weckesser, J., 1993. Cyanopeptolins, new depsipeptides from the cyanobacterium Microcystis sp. pcc 7806. J. Antibiot. (Tokyo) 46, 1550–1556. https://doi.org/10.7164/antibiotics.46.1550

McKindles, K., Frenken, T., McKay, R.M.L., Bullerjahn, G.S., 2020. Binational efforts addressing cyanobacterial harmful algal blooms in the great lakes. Handb. Environ. Chem. 101, 109–133. https://doi.org/10.1007/698_2020_513/FIGURES/3

Meyer, K.A., Davis, T.W., Watson, S.B., Denef, V.J., Berry, M.A., Dick, G.J., 2017. Genome sequences of lower Great Lakes Microcystis sp. reveal strain-specific genes that are present and expressed in western Lake Erie blooms. PLOS ONE 12, e0183859. https://doi.org/10.1371/journal.pone.0183859

Mikalsen, B., Boison, G., Skulberg, O.M., Fastner, J., Davies, W., Gabrielsen, T.M., Rudi, K., Jakobsen, K.S., 2003. Natural variation in the microcystin synthetase operon mcyABC and impact on microcystin production in Microcystis strains. J. Bacteriol. 185, 2774–85. https://doi.org/10.1128/jb.185.9.2774-2785.2003

Miller, T.R., Beversdorf, L.J., Weirich, C.A., Bartlett, S.L., 2017. Cyanobacterial Toxins of the Laurentian Great Lakes, Their Toxicological Effects, and Numerical Limits in Drinking Water. Mar. Drugs 15, 160. https://doi.org/10.3390/md15060160

Moreira, C., Spillane, C., Fathalli, A., Vasconcelos, V., Antunes, A., 2014. African origin and europe-mediated global dispersal of the cyanobacterium Microcystis aeruginosa. Curr. Microbiol. 69, 628–633. https://doi.org/10.1007/s00284-014-0628-2

Morris, J.J., Kirkegaard, R., Szul, M.J., Johnson, Z.I., Zinser, E.R., 2008. Facilitation of Robust Growth of Prochlorococcus Colonies and Dilute Liquid Cultures by “Helper” Heterotrophic Bacteria. Appl. Environ. Microbiol. 74, 4530–4534. https://doi.org/10.1128/AEM.02479-07

Mutsuda, M., Ishikawa, T., Takeda, T., Shigeoka, S., 1996. The catalase-peroxidase of Synechococcus PCC 7942: purification, nucleotide sequence analysis and expression in Escherichia coli. Biochem. J. 316, 251–257. https://doi.org/10.1042/bj3160251

Ogawa, T., Marco, E., Orus, M.I., 1994. A gene (ccmA) required for carboxysome formation in the cyanobacterium Synechocystis sp. strain PCC6803. J. Bacteriol. 176, 2374–2378. https://doi.org/10.1128/jb.176.8.2374-2378.1994

Palagama, D.S.W., Baliu-Rodriguez, D., Snyder, B.K., Thornburg, J.A., Bridgeman, T.B., Isailovic, D., 2020. Identification and quantification of microcystins in western Lake Erie during 2016 and 2017 harmful algal blooms. J. Gt. Lakes Res. 46, 289–301. https://doi.org/10.1016/j.jglr.2020.01.002

Parks, D.H., Chuvochina, M., Waite, D.W., Rinke, C., Skarshewski, A., Chaumeil, P.-A., Hugenholtz, P., 2018. A standardized bacterial taxonomy based on genome phylogeny substantially revises the tree of life. Nat Biotechnol. https://doi.org/10.1038/nbt.4229

Pawlik-Skowrońska, B., Bownik, A., 2022. Synergistic toxicity of some cyanobacterial oligopeptides to physiological activities of Daphnia magna (Crustacea). Toxicon 206, 74–84. https://doi.org/10.1016/j.toxicon.2021.12.013

Pérez-Carrascal, O.M., Terrat, Y., Giani, A., Fortin, N., Greer, C.W., Tromas, N., Shapiro, B.J., 2019. Coherence of Microcystis species revealed through population genomics. bioRxiv 541755. https://doi.org/10.1101/541755

Price, M.N., Dehal, P.S., Arkin, A.P., 2009. FastTree: Computing Large Minimum Evolution Trees with Profiles instead of a Distance Matrix. Mol. Biol. Evol. 26, 1641–1650. https://doi.org/10.1093/MOLBEV/MSP077

Puddick, J., Prinsep, M., Wood, S., Kaufononga, S., Cary, S., Hamilton, D., 2014. High Levels of Structural Diversity Observed in Microcystins from Microcystis CAWBG11 and Characterization of Six New Microcystin Congeners. Mar. Drugs 12, 5372–5395. https://doi.org/10.3390/md12115372

Raven, J.A., Gobler, C.J., Hansen, P.J., 2020. Dynamic CO2 and pH levels in coastal, estuarine, and inland waters: Theoretical and observed effects on harmful algal blooms. Harmful Algae, Climate change and harmful algal blooms 91, 101594. https://doi.org/10.1016/j.hal.2019.03.012

Rinta-Kanto, J., Ouellette, A., Boyer, G., Twiss, M., Bridgeman, T., Wilhelm, S., 2005. Quantification of Toxic Microcystis spp. during the 2003 and 2004 Blooms in Western Lake Erie using Quantitative Real-Time PCR. Environ. Sci. Technol. 39, 4198–4205. https://doi.org/10.1021/ES048249U

Rinta-Kanto, J.M., Konopko, E.A., DeBruyn, J.M., Bourbonniere, R.A., Boyer, G.L., Wilhelm, S.W., 2009. Lake Erie Microcystis: Relationship between microcystin production, dynamics of genotypes and environmental parameters in a large lake. Harmful Algae 8, 665–673. https://doi.org/10.1016/j.hal.2008.12.004

Rounge, T.B., Rohrlack, T., Tooming-Klunderud, A., Kristensen, T., Jakobsen, K.S., 2007. Comparison of cyanopeptolin genes in Planktothrix, Microcystis, and Anabaena strains: Evidence for independent evolution within each genus. Appl. Environ. Microbiol. 73, 7322–7330. https://doi.org/10.1128/AEM.01475-07

Sandrini, G., Cunsolo, S., Merijn Schuurmans, J., Matthijs, H.C.P., Huisman, J., 2015. Changes in gene expression, cell physiology and toxicity of the harmful cyanobacterium Microcystis aeruginosa at elevated CO2. Front. Microbiol. 6, 401. https://doi.org/10.3389/fmicb.2015.00401

Sandrini, G., Matthijs, H.C.P., Verspagen, J.M.H., Muyzer, G., Huisman, J., 2014. Genetic diversity of inorganic carbon uptake systems causes variation in CO2 response of the cyanobacterium Microcystis. ISME J. 8, 589–600. https://doi.org/10.1038/ismej.2013.179

Santos, A.A., Soldatou, S., de Magalhães, V.F., Azevedo, S.M.F.O., Camacho-Muñoz, D., Lawton, L.A., Edwards, C., 2021. Degradation of Multiple Peptides by Microcystin-Degrader Paucibacter toxinivorans (2C20). Toxins 13, 265. https://doi.org/10.3390/toxins13040265

Schloss, P.D., Handelsman, J., 2005. Metagenomics for studying unculturable microorganisms: Cutting the Gordian knot. Genome Biol. 6, 1–4. https://doi.org/10.1186/GB-2005-6-8-229/TABLES/1

Schuurmans, J.M., Brinkmann, B.W., Makower, A.K., Dittmann, E., Huisman, J., Matthijs, H.C.P., 2018. Microcystin interferes with defense against high oxidative stress in harmful cyanobacteria. Harmful Algae 78, 47–55. https://doi.org/10.1016/j.hal.2018.07.008

Seymour, J.R., Amin, S.A., Raina, J.B., Stocker, R., 2017. Zooming in on the phycosphere: The ecological interface for phytoplankton-bacteria relationships. Nat. Microbiol. 2, 1–12. https://doi.org/10.1038/nmicrobiol.2017.65

Shimizu, K., Maseda, H., Okano, K., Kurashima, T., Kawauchi, Y., Xue, Q., Utsumi, M., Zhang, Z., Sugiura, N., 2012. Enzymatic pathway for biodegrading microcystin LR in Sphingopyxis sp. C1. J. Biosci. Bioeng. 114, 630–634. https://doi.org/10.1016/j.jbiosc.2012.07.004

Smith, D.J., Berry, M.A., Cory, R.M., Johengen, T.H., Kling, G.W., Davis, T.W., Dick, G.J., 2022a. Heterotrophic Bacteria Dominate Catalase Expression during Microcystis Blooms. Appl. Environ. Microbiol. 88, e02544–21. https://doi.org/10.1128/aem.02544-21

Smith, D.J., Kharbush, J.J., Kersten, R.D., Dick, G.J., 2022b. Uptake of Phytoplankton-Derived Carbon and Cobalamins by Novel Acidobacteria Genera in Microcystis Blooms Inferred from Metagenomic and Metatranscriptomic Evidence. Appl. Environ. Microbiol. 88, e01803–21. https://doi.org/10.1128/aem.01803-21

Smith, D.J., Tan, J.Y., Powers, M.A., Lin, X.N., Davis, T.W., Dick, G.J., 2021. Individual Microcystis colonies harbour distinct bacterial communities that differ by Microcystis oligotype and with time. Environ. Microbiol. 23, 3020–3036. https://doi.org/10.1111/1462-2920.15514

Smith, R.B., Bass, B., Sawyer, D., Depew, D., Watson, S.B., 2019. Estimating the economic costs of algal blooms in the Canadian Lake Erie Basin. Harmful Algae 87, 101624. https://doi.org/10.1016/j.hal.2019.101624

Steffen, M.M., Davis, T.W., McKay, R.M.L., Bullerjahn, G.S., Krausfeldt, L.E., Stough, J.M.A., Neitzey, M.L., Gilbert, N.E., Boyer, G.L., Johengen, T.H., Gossiaux, D.C., Burtner, A.M., Palladino, D., Rowe, M.D., Dick, G.J., Meyer, K.A., Levy, S., Boone, B.E., Stumpf, R.P., Wynne, T.T., Zimba, P.V., Gutierrez, D., Wilhelm, S.W., 2017. Ecophysiological Examination of the Lake Erie *Microcystis* Bloom in 2014: Linkages between Biology and the Water Supply Shutdown of Toledo, OH. Environ. Sci. Technol. 51, 6745–6755. https://doi.org/10.1021/acs.est.7b00856

Stemberger, R.S., 1981. A General Approach to the Culture of Planktonic Rotifers. Can. J. Fish. Aquat. Sci. 38, 721–724. https://doi.org/10.1139/f81-095

Tang, Y.Z., Koch, F., Gobler, C.J., 2010. Most harmful algal bloom species are vitamin B1 and B12 auxotrophs. Proc. Natl. Acad. Sci. 107, 20756–20761. https://doi.org/10.1073/pnas.1009566107

Tillett, D., Dittmann, E., Erhard, M., Von Döhren, H., Börner, T., Neilan, B.A., 2000. Structural organization of microcystin biosynthesis in Microcystis aeruginosa PCC7806: An integrated peptide-polyketide synthetase system. Chem. Biol. 7, 753–764. https://doi.org/10.1016/S1074-5521(00)00021-1

Tonk, L., Van De Waal, D.B., Slot, P., Huisman, J., Matthijs, H.C.P., Visser, P.M., 2008. Amino acid availability determines the ratio of microcystin variants in the cyanobacterium Planktothrix agardhii. FEMS Microbiol. Ecol. 65, 383–390. https://doi.org/10.1111/j.1574-6941.2008.00541.x

Tooming-Klunderud, A., Mikalsen, B., Kristensen, T., Jakobsen, K.S., 2008. The mosaic structure of the mcyABC operon in Microcystis. Microbiology 154, 1886–1899. https://doi.org/10.1099/mic.0.2007/015875-0

US EPA, O., 2019. Method 546: Determination of Total Microcystins and Nodularins in Drinking Water and Ambient Water by Adda Enzyme-Linked Immunosorbent Assay [WWW Document]. URL https://www.epa.gov/esam/method-546-determination-total-microcystins-and-nodularins-drinking-water-and-ambient-water (accessed 9.15.22).

Wagner, N.D., Osburn, F.S., Wang, J., Taylor, R.B., Boedecker, A.R., Chambliss, C.K., Brooks, B.W., Scott, J.T., 2019. Biological stoichiometry regulates toxin production in microcystis aeruginosa (UTEX 2385). Toxins 11, 601. https://doi.org/10.3390/toxins11100601

Watson, S.B., Miller, C., Arhonditsis, G., Boyer, G.L., Carmichael, W., Charlton, M.N., Confesor, R., Depew, D.C., Höök, T.O., Ludsin, S.A., Matisoff, G., McElmurry, S.P., Murray, M.W., Peter Richards, R., Rao, Y.R., Steffen, M.M., Wilhelm, S.W., 2016. The re-eutrophication of Lake Erie: Harmful algal blooms and hypoxia. Harmful Algae 56, 44–66. https://doi.org/10.1016/J.HAL.2016.04.010

Wood, D.E., Lu, J., Langmead, B., 2019. Improved metagenomic analysis with Kraken 2. Genome Biol 20, 257. https://doi.org/10.1186/s13059-019-1891-0

Wood, D.E., Salzberg, S.L., 2014. Kraken: ultrafast metagenomic sequence classification using exact alignments. Genome Biol. 15, R46. https://doi.org/10.1186/gb-2014-15-3-r46

Woodcroft, B.J., n.d. CoverM: Read coverage calculator for metagenomics.

Yancey, C.E., Smith, D.J., Uyl, P.A.D., Mohamed, O.G., Yu, F., Ruberg, S.A., Chaffin, J.D., Goodwin, K.D., Tripathi, A., Sherman, D.H., Dick, G.J., 2022a. Metagenomic and Metatranscriptomic Insights into Population Diversity of Microcystis Blooms: Spatial and Temporal Dynamics of mcy Genotypes, Including a Partial Operon That Can Be Abundant and Expressed. Appl. Environ. Microbiol. https://doi.org/10.1128/AEM.02464-21

Yancey, C.E., Yu, F., Tripathi, A., Sherman, D.H., Dick, G.J., 2022b. Expression of Microcystis biosynthetic gene clusters in natural populations suggests temporally dynamic synthesis of novel and known secondary metabolites in western Lake Erie. https://doi.org/10.1101/2022.10.12.511943

Youngblut, N.D., Ley, R.E., 2021. Struo2: efficient metagenome profiling database construction for ever-expanding microbial genome datasets. PeerJ 9, e12198. https://doi.org/10.7717/peerj.12198

