## Supplementary material for "The Western Lake Erie Culture Collection: A promising resource for evaluating the physiological and genetic diversity of *Microcystis* and its associated microbiome": Table S

**Supplemental Information**

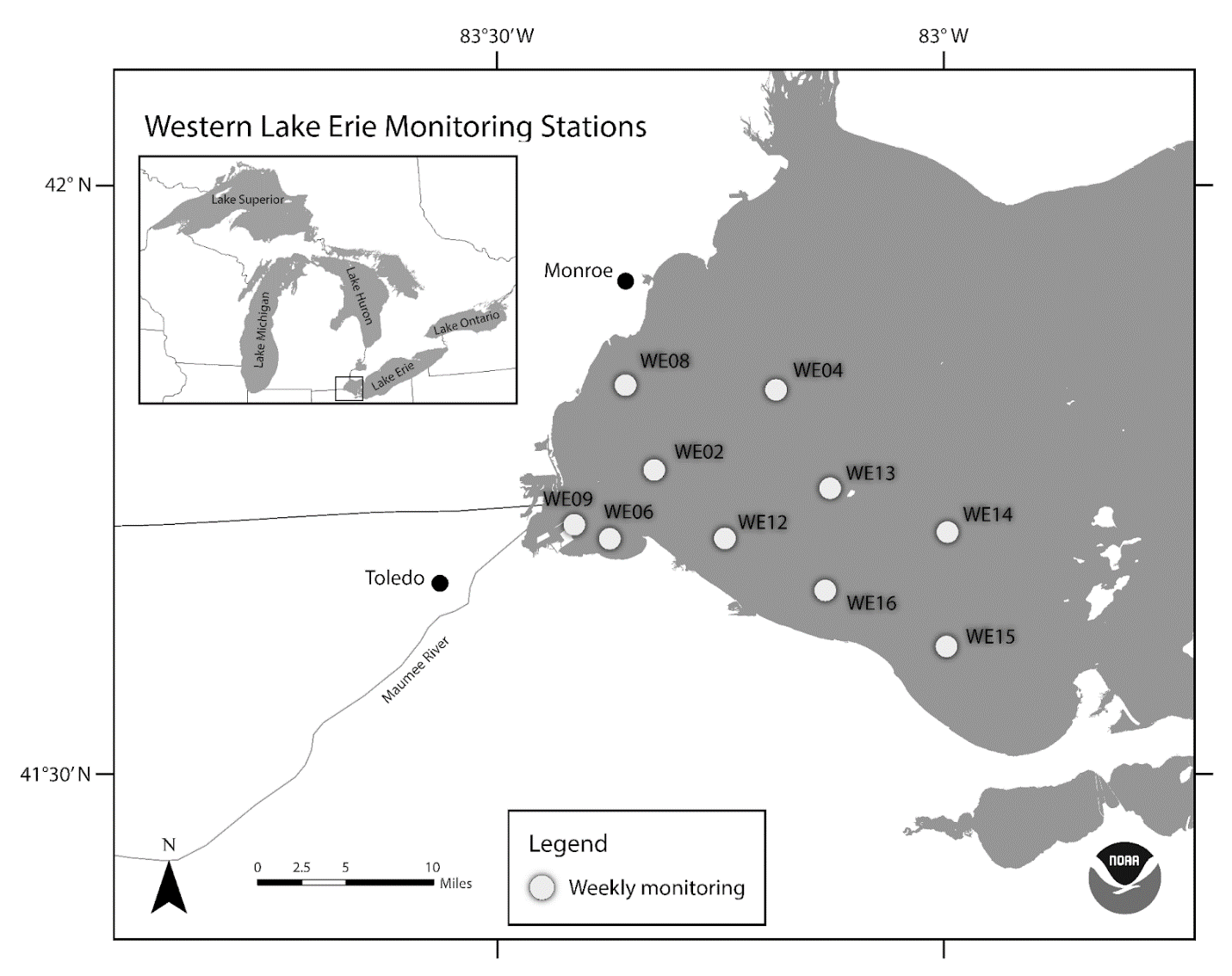

Figure S1: Western Lake Erie cyanoHAB monitoring stations as part of the long term monitoring efforts by the NOAA Great Lakes Environmental Research Laboratory (GLERL), and Cooperative Institute for Great Lakes Research (CIGLR). WLECC isolates were collected and isolated from WE2, WE4, WE6, WE8, and WE12.

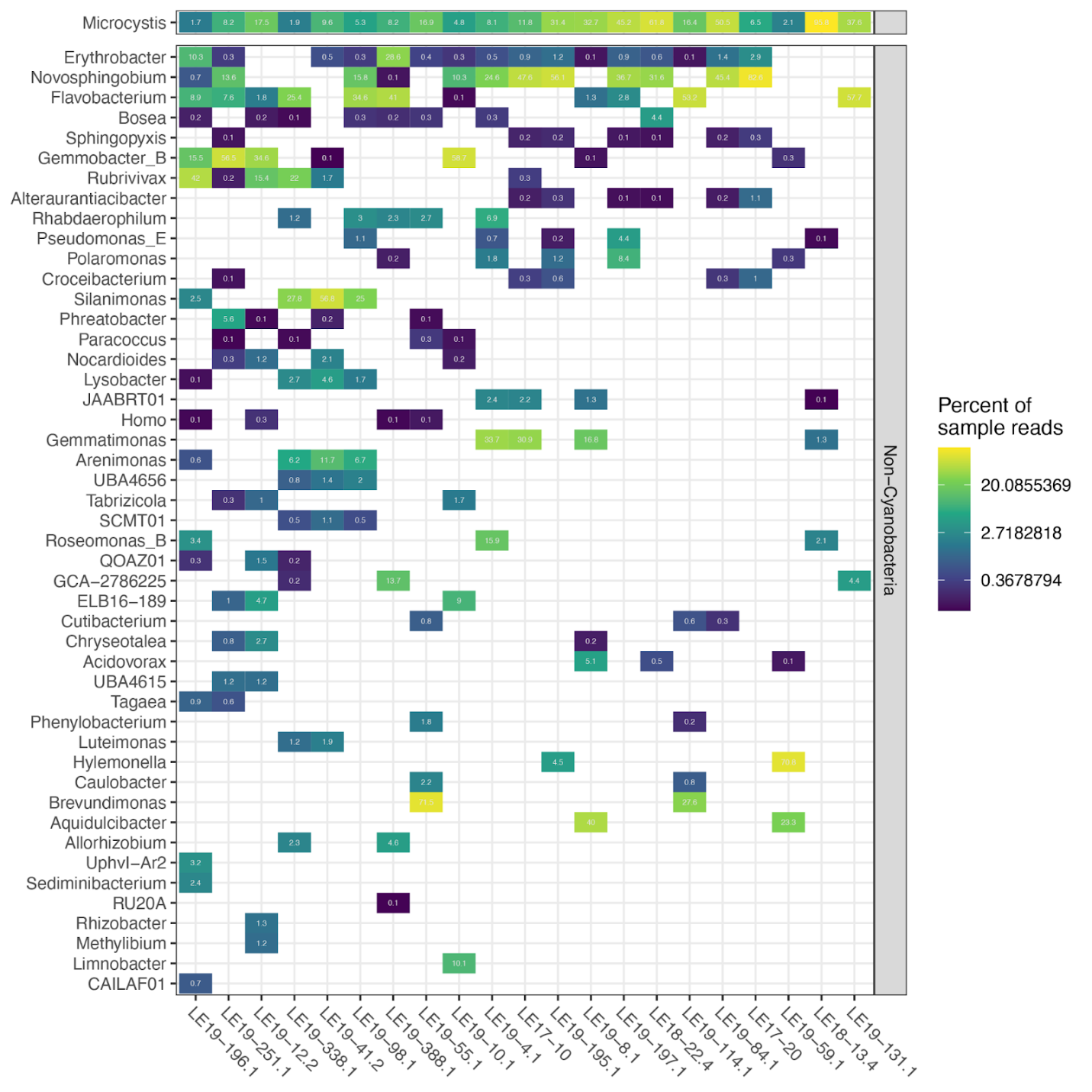

Figure S2: Culture composition by read profiling with Kraken and Bracken

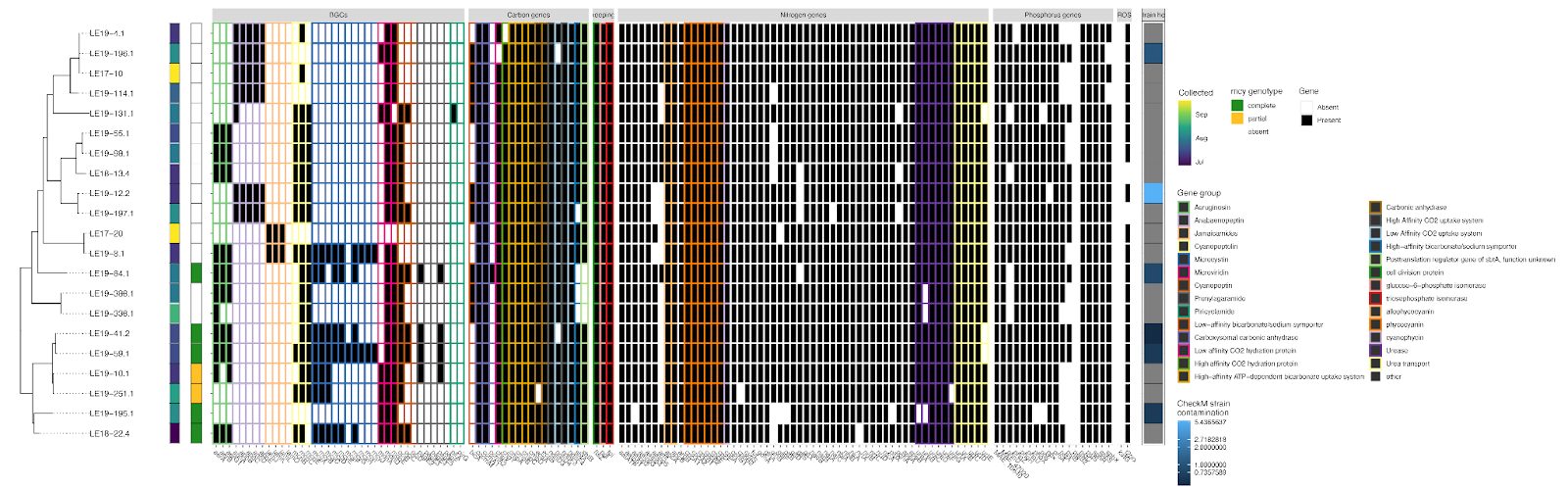

Figure S3: WLE *Microcystis* strains gene content without removal of fully present and absent genes.

Table S1: List of Genes used to query *Microcystis* and associated bacteria genomes

|  | **Gene Symbol** | **NCBI Accession** | **Function** |
| --- | --- | --- | --- |
| Secondary Metabolism | apn | KU665241.1 | anabenopeptin synthesis |
|  | sxt | DQ787200.1 | saxitoxin synthesis |
|  | ana | JF803645.1 | anatoxin synthesis |
|  | nda | HQ682080.1 | nodularin synthesis |
|  | cyr | EU140798.1 | cylindrospermopsin synthesis |
|  | jam | AY522504.1 | jamaicamide synthesis |
|  | ltx | AY588942.1 | lyngbyatoxin synthesis |
|  | pag | HQ655154.1 | prenylagaramide synthesis |
|  | oci | KU665241.1 | cyanopeptin synthesis |
|  | mcy | AF183408.1 | microcystin synthesis |
|  | aer | FJ609416.1. | aeruginosin synthesis |
|  | mcn | DQ075244.1. | cyanopeptolin synthesis |
|  | pir | JQ951924.1. | piricyclamide synthesis |
|  | mdn | AM943877.1. | microviridin synthesis |
| P Metabolism | ppk | AP009552.1 | polyphosphate kinase |
|  | pstB | AP009552.1 | phosphate ABC transporter ATP-binding protein |
|  | pstA | AP009552.1 | phosphate transport system permease protein |
|  | pstC | AP009552.1 | phosphate transport system permease protein |
|  | pstC | AP009552.1 | ABC-transporter periplasmic phosphate binding protein |
|  | phnD | AP009552.1 | ABC-transporter substrate binding protein |
|  | akp | AP009552.1 | alkaline phosphatase |
|  | pstA | AP009552.1 | phosphate transport system permease protein |
|  | pstA | AP009552.1 | phosphate transport system permease protein |
|  | pstA | AP009552.1 | phosphate transport system permease protein |
|  | pstS | AP009552.1 | phosphate-binding periplasmic protein |
|  | pstB2 | AP009552.1 | phosphate transport ATP-binding protein |
|  | pstB | AP009552.1 | phosphate transport ATP-binding protein |
|  | pstA | AP009552.1 | phosphate ABC transporter permease |
|  | pstC | AP009552.1 | phosphate ABC transporter permease |
|  | pstS | AP009552.1 | phosphate-binding periplasmic protein |
|  | sphX | AP009552.1 | phosphate transport system substrate binding protein |
|  | phoX | AP009552.1 | alkaline phosphatase |
|  | proA | AP009552.1 | gamma-glutamyl phosphate reductase |
|  | pstS | AP009552.1 | phosphate binding protein PstS homolog |
|  | phoH | AP009552.1 | phoH like protein |
|  | sipp | AP009552.1 | soluble inorganic pyrophosphatase |
|  | alp2 | AP009552.1 | alkaline phosphatase-like protein |
|  | phnZ | AP009552.1 | metal dependent phosphohyrolase HD region |
|  | sphR | AP009552.1 | response regulator in two component regulatory system of Pi uptake |
|  | sphS | AP009552.1 | two-component sensor histidine kinase |
|  | phoU | AP009552.1 | phosphate transport system regulatory protein |
|  | ppx | AP009552.1 | exopolyphosphatase |
| N metabolism | natC | CP020771.1 | neutral amino acid ABC transporter membrane protein |
|  | natA | CP020771.1 | neutral amino acid ABC transporter ATP-binding protein |
|  | ntcA | CP020771.1 | Global N regulator |
|  | nirA | CP020771.1 | Ferridoxin-nitrate reductase |
|  | ureD | CP020771.1 | Urease accessory protein D |
|  | urtE | CP020771.1 | Urea transport |
|  | urtD | CP020771.1 | Urea transport |
|  | urtC | CP020771.1 | Urea transport |
|  | urtB | CP020771.1 | Urea transport |
|  | urtA | CP020771.1 | Urea transport |
|  | gltB | CP020771.1 | Glutamate synthase (ferridoxin) |
|  | gdhA | CP020771.1 | Glutamate dehydrogenase (NADP) |
|  | glnA | CP020771.1 | L-glutamine synthetase |
|  | cynS | CP020771.1 | Cyanate lyase |
|  | amt | CP020771.1 | Ammonium transporter |
|  | gltS | CP020771.1 | monocomponent sodium-dependent glutamate permease |
|  | nrtD | CP020771.1 | Nitrate transport |
|  | nrtC | CP020771.1 | Nitrate transport |
|  | nrtB | CP020771.1 | Nitrate transport |
|  | nrtA | CP020771.1 | Nitrate transport |
|  | gltD | CP020771.1 | glutamate synthase (NADH) small subunit |
|  | speB | CP020771.1 | Urea metabolism (agmatinase) |
|  | ureG | CP020771.1 | urease accessory protein G |
|  | aatpp1 | CP020771.1 | Amino acid transport |
|  | aatATO | CP020771.1 | Amino acid transport |
|  | aatpp2 | CP020771.1 | Amino acid transport |
|  | aatpp3 | CP020771.1 | Amino acid transport |
|  | aatpp4 | CP020771.1 | Amino acid transport |
|  | aatpp5 | CP020771.1 | Amino acid transport |
|  | glsF | CP020771.1 | glutamate synthase (ferredoxin) |
|  | nadB | CP020771.1 | L-aspartate oxidase |
|  | fur | CP020771.1 | Ferric uptake regulation protein |
|  | ureF | CP020771.1 | urease |
|  | ureA | CP020771.1 | urease |
|  | ureB | CP020771.1 | urease |
|  | speA | CP020771.1 | Urea metabolism (agmatinase) |
|  | gltX | CP020771.1 | glutamyl-tRNA synthetase |
|  | narB | CP020771.1 | assimilatory nitrate reductase (ferredoxin) precursor |
|  | pipX | CP020771.1 | Nitrogen regulation |
|  | glnB | CP020771.1 | nitrogen regulatory protein P-II family |
|  | ureC | CP020771.1 | urease subunit alpha |
| N metabolism 2 (Laura) | cpcB1 | AP009552.1 | phycocyanin beta subunit |
|  | cpcB2 | AP009552.1 | phycocyanin beta subunit |
|  | cpcA1 | AP009552.1 | phycocyanin alpha subunit |
|  | cpcA2 | AP009552.1 | phycocyanin alpha subunit |
|  | cpcF | AP009552.1 | phycocyanin alpha-subunit phycocyanobilin lyase |
|  | cpcE | AP009552.1 | phycocyanin alpha-subunit phycocyanobilin lyase |
|  | nblA | AP009552.1 | phycobilisome degradation protein |
|  | nblB | AP009552.1 | phycocyanin alpha phycocyanobilin lyase related protein |
|  | argB | AP009552.1 | N-acetylglutamate kinase |
|  | cphB | AP009552.1 | cyanophycinase |
|  | cphA | AP009552.1 | cyanophycin synthetase |
|  | apcB | AP009552.1 | allophycocyanin beta subunit |
|  | apcA | AP009552.1 | allophycocyanin alpha subunit |
|  | pphA | AP009552.1 | protein serin/threonin phosphatase |
| CCM genes | ndhF3 | AP009552.1 | High Affinity CO2 uptake system |
|  | ndhD3 | AP009552.1 | High Affinity CO2 uptake system |
|  | ndhF4 | AP009552.1 | Low Affinity CO2 uptake system |
|  | ndhD4 | AP009552.1 | Low affinity CO2 uptake system subunit M |
|  | bicA | AP009552.1 | Low-affinity bicarbonate/sodium symporter |
|  | sbtA | AP009552.1 | High-affinity bicarbonate/sodium symporter |
|  | cmpA | AP009552.1 | High-affinity ATP-dependent bicarbonate uptake system |
|  | cmpB | AP009552.1 | High-affinity ATP-dependent bicarbonate uptake system |
|  | cmpC | AP009552.1 | High-affinity ATP-dependent bicarbonate uptake system |
|  | cmpD | AP009552.1 | High-affinity ATP-dependent bicarbonate uptake system |
|  | ecaA | AP009552.1 | Alpha carbonic anhydrase |
|  | ecaB | AP009552.1 | Beta carbonic anhydrase |
|  | ccmA1 | AP009552.1 | Carboxysomal carbonic anhydrase |
|  | ccmA2 | AP009552.1 | Carboxysomal carbonic anhydrase |
|  | cccmM | AP009552.1 | Carboxysomal carbonic anhydrase |
|  | sbtB | AP009552.1 | Posttranslation regulator gene of sbtA, function unknown |
|  | chpY | AP009552.1 | High affinity CO2 hydration protein |
|  | chpX | AP009552.1 | Low affinity CO2 hydration protein |
| Housekeeping | ftsZ | AP009552.1 | cell division protein |
|  | pgi | AP009552.1 | glucose-6-phosphate isomerase |
|  | tpi | AP009552.1 | triosephosphate isomerase |
| mlr (MC degradation) | mlrA | HMMs (see methods) | peptidase, hydrolyses microcystin LR to linear microcystin LR. |
|  | mlrB | HMMs (see methods) | active site serine peptidase, hydrolyses linear microcystin LR to tetrapeptide product |
|  | mlrC | HMMs (see methods) | peptidase, part of microcystin LR degradation pathway |
|  | mlrD | HMMs (see methods) | putative transporter/membrane anchoring |
| ROS | katE | JGI IMG genes with Kegg K03781 annotations | Catalase |
|  | katG | JGI IMG genes with Kegg K03782 annotations | Catalase-peroxidase |
|  | AHP1 | CAO86518.1 | t2prx |
|  | AHP1 | BAG06100.1 | t2prx |

Supplemental Table 2: Physicochemical measurements in at WLE stations during sample collection for culture isolates

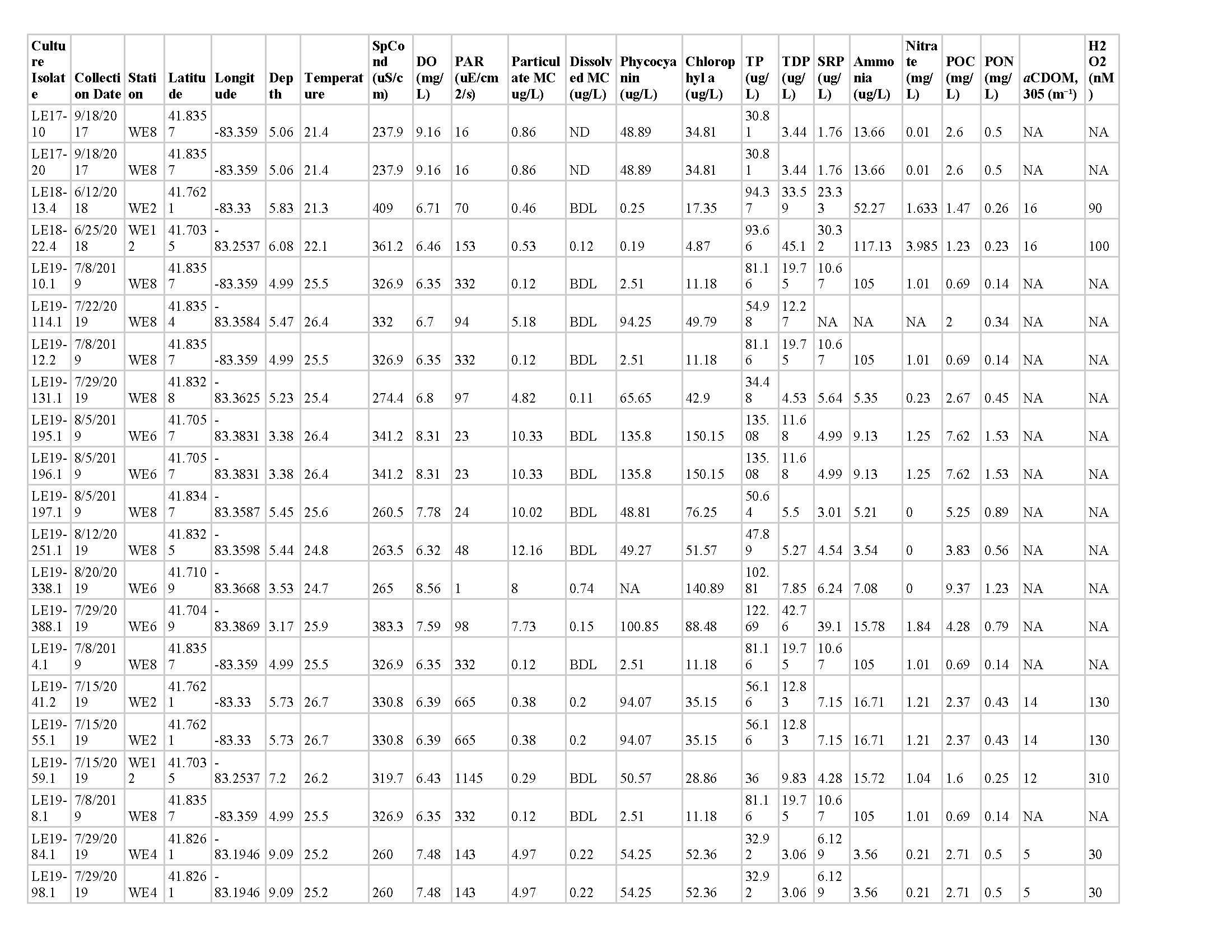

Supplemental Table 3: Bin statistics for *Microcystis* MAGs/ Completion, Contamination, and Strain Heterogeneity are measured using CheckM

| **Isolate** | **Completion** | **Contamination** | **Strain Het.** | **Strain contamination** | **N50** | **# of Contigs** |
| --- | --- | --- | --- | --- | --- | --- |
| LE19-4.1 | 99.67 | 0.73 | 0 | 0 | 41894 | 210 |
| LE17-10 | 99.48 | 0 | 0 | 0 | 41130 | 203 |
| LE19-13.4 | 99.89 | 0.07 | 0 | 0 | 39687 | 218 |
| LE19-55.1 | 99.89 | 0.07 | 0 | 0 | 39459 | 215 |
| LE19-98.1 | 99.89 | 0.07 | 0 | 0 | 37398 | 233 |
| LE19-8.1 | 99.89 | 0.29 | 0 | 0 | 32578 | 256 |
| LE17-20 | 99.89 | 0.07 | 0 | 0 | 31341 | 283 |
| LE19-197.1 | 99.89 | 0.29 | 0 | 0 | 28714 | 329 |
| LE19-131.1 | 94.63 | 0.28 | 0 | 0 | 26591 | 322 |
| LE19-114.1 | 99.89 | 0.29 | 0 | 0 | 25589 | 323 |
| LE19-388.1 | 99.89 | 0.4 | 0 | 0 | 20964 | 461 |
| LE19-84.1 | 99.48 | 0.39 | 50 | 0.20 | 19776 | 532 |
| LE19-195.1 | 89.14 | 0.95 | 87.5 | 0.83 | 16572 | 362 |
| LE18-22.4 | 99.89 | 0.15 | 0 | 0 | 14457 | 642 |
| LE19-251.1 | 96.38 | 0.07 | 0 | 0 | 12280 | 643 |
| LE19-10.1 | 99.45 | 0.07 | 0 | 0 | 11313 | 772 |
| LE19-338.1 | 98.94 | 0.4 | 0 | 0 | 9020 | 789 |
| LE19-59.1 | 98.79 | 1.29 | 44.44 | 0.57 | 8519 | 887 |
| LE19-12.2 | 98.46 | 7.07 | 81.40 | 5.75 | 8402 | 960 |
| LE19-196.1 | 94.08 | 3.59 | 30 | 1.1 | 4062 | 1365 |
| LE19-41.2 | 99.45 | 1.17 | 50 | 0.6 |  |  |

Supplemental Table 4: Read counts for *mcy* genes from *Microcystis* genomes

| Culture | *mcy* Genotype Status | *mcy* Gene | | | | | | | | | |
| --- | --- | --- | --- | --- | --- | --- | --- | --- | --- | --- | --- |
|  |  | mcyA | mcyB | mcyC | mcyD | mcyE | mcyF | mcyG | mcyH | mcyI | mcyJ |
| LE19-41.2 | Complete | 943 | 697 | 452 | 1512 | 1386 | 86 | 977 | 219 | 133 | 176 |
| LE18-22.4 | Complete | 9057 | 6684 | 4337 | 13589 | 12294 | 879 | 8398 | 1807 | 1330 | 1572 |
| LE19-281.1 | Complete | 294 | 200 | 122 | 415 | 398 | 23 | 254 | 51 | 52 | 48 |
| LE19-84.1 | Complete | 10134 | 7885 | 3593 | 14209 | 12440 | 951 | 9447 | 2011 | 1332 | 1664 |
| LE19-195.1 | Complete | 6984 | 5912 | 3134 | 10043 | 9676 | 710 | 6807 | 1525 | 1155 | 1280 |
| LE19-59.1 | Complete | 482 | 357 | 241 | 720 | 657 | 42 | 466 | 105 | 100 | 87 |
| LE19-10.1 | Partial | 303 | 741 | 444 | 0 | 0 | 0 | 0 | 0 | 0 | 0 |
| LE19-251.1 | Partial | 547 | 1254 | 748 | 0 | 0 | 0 | 0 | 0 | 0 | 0 |
| LE19-12.2 | Absent | 0 | 0 | 0 | 0 | 0 | 0 | 0 | 0 | 0 | 0 |
| LE17-20.1 | Absent | 1 | 0 | 0 | 0 | 0 | 0 | 2 | 0 | 0 | 0 |
| LE17-10 | Absent | 0 | 0 | 0 | 0 | 0 | 0 | 1 | 0 | 0 | 0 |
| LE19-114.1 | Absent | 0 | 4 | 2 | 2 | 0 | 0 | 1 | 0 | 4 | 0 |
| LE19-13.4 | Absent | 0 | 0 | 0 | 0 | 0 | 0 | 0 | 0 | 0 | 0 |
| LE19-196.1 | Absent | 0 | 0 | 0 | 0 | 0 | 0 | 0 | 0 | 0 | 0 |
| LE19-338.1 | Absent | 1 | 0 | 0 | 0 | 0 | 0 | 0 | 0 | 0 | 0 |
| LE19-55.1 | Absent | 0 | 0 | 0 | 0 | 0 | 0 | 0 | 0 | 0 | 0 |
| LE19-8.1 | Absent | 0 | 3 | 0 | 0 | 2 | 0 | 2 | 0 | 0 | 0 |
| LE19-131.1 | Absent | 0 | 0 | 0 | 0 | 0 | 0 | 0 | 0 | 0 | 0 |
| LE19-197.1 | Absent | 0 | 0 | 0 | 0 | 0 | 0 | 0 | 0 | 0 | 0 |
| LE19-4.1 | Absent | 0 | 2 | 0 | 0 | 0 | 0 | 0 | 0 | 0 | 0 |
| LE19-98.1 | Absent | 0 | 0 | 0 | 6 | 2 | 0 | 4 | 0 | 0 | 0 |
| LE17-20 | Absent | 0 | 0 | 0 | 0 | 0 | 0 | 0 | 0 | 0 | 0 |
| LE19-388.1 | Absent | 2 | 0 | 0 | 0 | 0 | 0 | 0 | 0 | 0 | 0 |
